# Spatial transcriptomic characterization of COVID-19 pneumonitis identifies immune circuits related to tissue injury

**DOI:** 10.1101/2021.06.21.449178

**Authors:** AR Cross, CE de Andrea, Villalba-Esparza María, MF Landecho Acha, L Cerundolo, P Weeratunga, R Etherington, L Denney, G Ogg, LP Ho, ISD Roberts, J Hester, P Klenerman, I Melero, SN Sansom, F Issa

## Abstract

Severe lung damage in COVID-19 involves complex interactions between diverse populations of immune and stromal cells. In this study, we used a spatial transcriptomics approach to delineate the cells, pathways and genes present across the spectrum of histopathological damage in COVID-19 lung tissue. We applied correlation network-based approaches to deconvolve gene expression data from areas of interest within well preserved post-mortem lung samples from three patients. Despite substantial inter-patient heterogeneity we discovered evidence for a common immune cell signaling circuit in areas of severe tissue that involves crosstalk between cytotoxic lymphocytes and pro-inflammatory macrophages. Expression of *IFNG* by cytotoxic lymphocytes was associated with induction of chemokines including *CXCL9, CXCL10* and *CXCL11* which are known to promote the recruitment of CXCR3+ immune cells. The tumour necrosis factor (TNF) superfamily members *BAFF* (*TNFSF13B*) and *TRAIL* (*TNFSF10*) were found to be consistently upregulated in the areas with severe tissue damage. We used published spatial and single cell SARS-CoV-2 datasets to confirm our findings in the lung tissue from additional cohorts of COVID-19 patients. The resulting model of severe COVID-19 immune-mediated tissue pathology may inform future therapeutic strategies.

**One Sentence Summary:** Spatial analysis identifies IFNγ response signatures as focal to severe alveolar damage in COVID-19 pneumonitis.

## INTRODUCTION

Mortality following severe acute respiratory syndrome coronavirus 2 (SARS-CoV-2) infection is largely related to the antiviral response and immune-mediated lung injury (*1*). Histopathologically, coronavirus disease 2019 (COVID-19) pneumonitis is associated with diffuse alveolar damage (DAD), fibrosis, leukocytic infiltrates, and microvascular thromboses (*2–4*). Features of DAD include alveolar wall thickening, interstitial expansion, hyaline membrane deposition, and pneumocyte hyperplasia. Studies have begun to describe the transcriptomic profiles of lung pathology, although these have been largely designed to assess the cellular impact of SARS-CoV-2 infection (*5–7*). To the best of our knowledge, later-stage severe organ pathology is not consistently associated with high levels of infection or active viral replication (*8, 9*). In lung tissue from severe cases, the variability in the detection of SARS-CoV-2 RNA or antigen supports a model of inflammation-perpetuated disease (*5, 9*). The immune contributors and biological pathways associated with the widespread severe alveolar injury remain unclear, therefore a greater understanding of the pathological features of COVID-19 would complement the growing knowledge of both tissue and blood-based immune profiles (*10*).

Advanced spatial profiling techniques provide the tools to identify the distribution of proteins and RNAs *in situ*, allowing the dissection of biological processes in and around specific histological features of interest (*11, 12*). We employed an advanced multiplexed *in situ* hybridization tissue analysis platform to generate detailed transcriptomic profiles of multiple spatially discrete areas in lung samples from three COVID-19 patients with a focus on the spectrum of DAD severity. Uniquely, these tissues were obtained via open sampling at the point of death, which ensured high quality RNA analyses and avoided the caveats associated with late autopsies. Application of network-based approaches allowed for deconvolution and visualisation of the cell types and immune cell signaling phenotypes present in the patient tissues. Following integration of our results with those from other spatial and single cell sequencing studies, we propose a cellular model for the active immune processes in the lung during severe COVID-19.

## RESULTS

### Subhead 1: Immune cell infiltration is associated with severe local tissue damage in COVID-19

The histological and immune cell landscape within COVID-19 lung tissue from three patients with fatal disease was investigated to establish the extent of intra-tissue variation in cellular pathology (mean specimen area 1.78cm^2^; **Supplementary Table 1** for patient information). Each sample featured a spectrum of DAD, from mild to severe, comprising pneumocyte hyperplasia, hyaline membrane formation, and interstitial expansion. There was a non-uniform distribution of immune infiltrates (**Figure 1A-B and Supplementary Table 2**). Histological features beyond DAD included vacuolated macrophages, oedema, vascular thrombi, and squamous metaplasia. Bulk detection for SARS-CoV-2 nucleocapsid RNA by qPCR in patient lung samples found low viral loads (**Supplementary Table 2**), whilst there was significant nucleocapsid protein within hyaline membranes and pneumocytes in patient B (**Supplementary Figure 1B and 1D**). By contrast, in patients A and C, only small areas of weak viral nucleocapsid protein expression were detected in the alveolar space and ciliated bronchiolar epithelium (**Supplementary Figure 1A, 1C-D**). Immunofluorescent staining for CD3, CD68 and pan-cytokeratin (panCK) enabled the identification of lymphocytes, macrophages, epithelial cells, and general tissue architecture (**Figure 1B**). Based on this staining we selected 46 areas of interest (AOIs) with a broad range of inflammatory features (**Figure 1B and 1C**). DAD histopathological severity was assessed by two independent pathologists and each AOI categorised as having either mild/moderate injury with some conservation of alveolar architecture or severe injury with a loss of alveolar structure and significant inflammation. In addition, for each AOI we quantified the number of CD3+ lymphocytes, CD68+ macrophages and cells (by nuclear stain). The SARS-CoV-2 nucleocapsid (N) protein positivity of adjacent tissue from sequential sections was also recorded. Areas of interest with severe DAD were associated with higher frequencies of T cells, although CD68+ macrophage numbers were not consistently increased (**Figure 1C and Supplementary Figure 1G-H**), but histopathological severity was not consistently associated with higher levels of RNA for the SARS-CoV-2 N protein (**Supplementary Figure 1E**).

**Figure 1:**
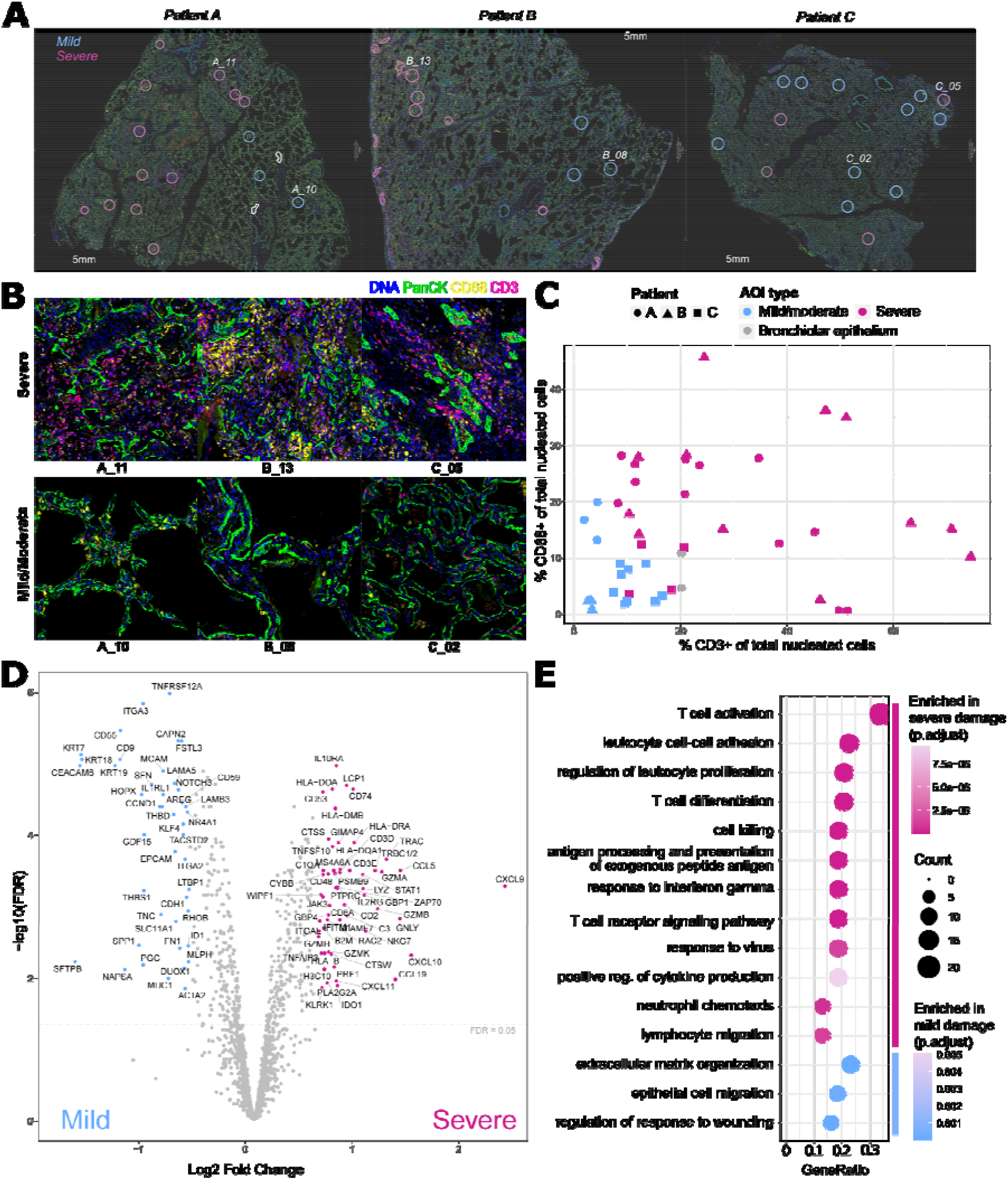
A spectrum of diffuse alveolar damage and inflammation was observed within and across COVID-19 lung biopsies. Selection and annotation of areas of diffuse alveolar damage in COVID-19 lung tissues for transcriptomic analysis. (**A**) Merged immunofluorescence (IF) images of the lung samples from patients A, B and C (scale bars: 5mm, pan cytokeratin-green, DNA-blue, CD3-red, CD68-yellow). n=47 areas of interest (AOI) selected for transcript profiling are highlighted (blue circles-mild/moderate and magenta circles-severe pathology), of which n=46 passed quality control after sequencing. Labelled areas correspond to higher magnification examples in **B**. (**B**) Representative IF images of AOIs demonstrating the morphology and immune infiltrate observed within areas of severe and mild/moderate diffuse alveolar damage. AOIs spanned on average 0.2mm^2^ (range: 0.05 to 0.33mm^2^) with exclusion of empty space. (**C**) The proportion of CD3^+^ and CD68^+^ cells of total nucleated cells were derived from the immunofluorescent imaging, plotted for each AOI and coloured by histological severity (mild/moderate-blue, severe-magenta) or inclusion of bronchiolar epithelium (grey). AOI are annotated by patient: A – circle, B – triangle and C - square. (**D**) Differential gene expression between areas of severe and mild/moderate damage. Coloured and annotated genes have a fold change expression > 1.5 and a Benjamini-Hochberg (BH) adjusted *p* value <0.05. (**E**) Selected GO biological processes significantly over-represented in genes differentially expressed between mild and severe areas of damage (BH corrected p < 0.05, one-sided Fisher’s exact test); see also **Supplementary Table 3** for differentially expressed genes and all over-represented pathways.

To profile the pathological pathways active in each AOI, we applied a multiplexed *in situ* hybridization approach to quantitate the expression of a curated panel of >1800 genes enriched in immune targets and augmented with a COVID-19-specific gene set. Use of robust quantile normalisation resulted in comparable distributions of gene expression and levels of housekeeping genes across the heterogenous AOIs (**Supplementary Figure 2**). Differential expression analysis of mild and severe DAD across all sampled AOIs (n=16 mild and n=28 severe) identified n=56 genes with significantly higher expression in severe pathology (Benjamini Hochberg [BH] adjusted p<0.05, |fold change| >1.5) including those encoding chemokines (*CXCL9, CXCL10, CXCL11, CCL19* and *CCL5*), cytotoxic molecules (*GZMA, GZMB, GZMK, PRF1, GNLY, LYZ, NKG7, KLRK1*), complement factors (*C1QA, C1QB*), and proteins involved in antigen processing and presentation (*CD74, HLA* genes, *CTSS*) (**Figure 1D**). Genes upregulated in the severe areas showed a significant over-representation (BH adjusted p<0.05) of gene ontology (GO) biological process terms related to T cell activation and differentiation, antigen presentation, cytokine production, cytotoxicity, and response to interferon gamma (**Figure 1E**). By contrast, genes enriched in areas of mild damage (n=40, BH adjusted p<0.05, |fold change| >1.5) had a significant over-representation (BH adjusted p<0.05) of pathways associated with wound healing, epithelial migration and extracellular matrix organisation (**Figure 1D**).

### Subhead 2: Network analysis implicates CD8^+^ T cells, mononuclear phagocytes and active TLR, interferon and IL-1 signaling in COVID-19 lung inflammation

To perform an unbiased exploration of the cellular and phenotypic variation present in the set of the profiled AOIs, we employed weighted gene correlation network analysis (WGCNA) (*13*). This analysis identified 17 distinct modules of co-expressed genes (n=27-266 genes per module, median 88) (**Supplementary Figure 3A-D and Supplementary Table 4**). We began a systematic characterization of the identified modules by investigating their association with specific cell types. To do so we correlated expression of the module eigengenes (the representative module expression patterns) with separate cell type abundance estimates that were determined for each AOI by automatic cell type deconvolution analyses (*14*) (**Figure 2A and Supplementary Figure 4A-D**). To confirm the involvement of specific cell types we examined the correspondence of the cell type abundance predictions with the expression of a curated set of known cell type marker genes. While we generally saw a good agreement between the marker genes and the automatic predictions of cell type abundance, correlated expression of *MPO, ELANE* and *CTSG* suggested that neutrophils may be present in some of the AOIs (**Supplementary Figure 5A-B**). To help resolve this discrepancy we applied mass-cytometry imaging to sequential sections and examined the areas aligned with selected severe AOIs. This analysis revealed the presence of a substantial number of CD15^+^ neutrophils in an area of severe damage from patient B, whose tissues also showed strong staining for the SARS-CoV-2 N protein. Additionally, this analysis confirmed the expected presence of CD68^+^ macrophages and CD8^+^ T cells within severe areas, with only a small number of CD4^+^ cells being detected (**Supplementary Figure 6**). Next, we identified biological processes and pathways over-represented amongst the gene members of each WGCNA module (**Figure 2B and Supplementary Table 5**) and explored the correlation of the module eigengenes with genes involved in immune signaling, the complement system, viral infection and fibrosis (**Figure 2C)**. Based on these analyses we named each module according to its cell type associations and biological pathway enrichments.

**Figure 2:**
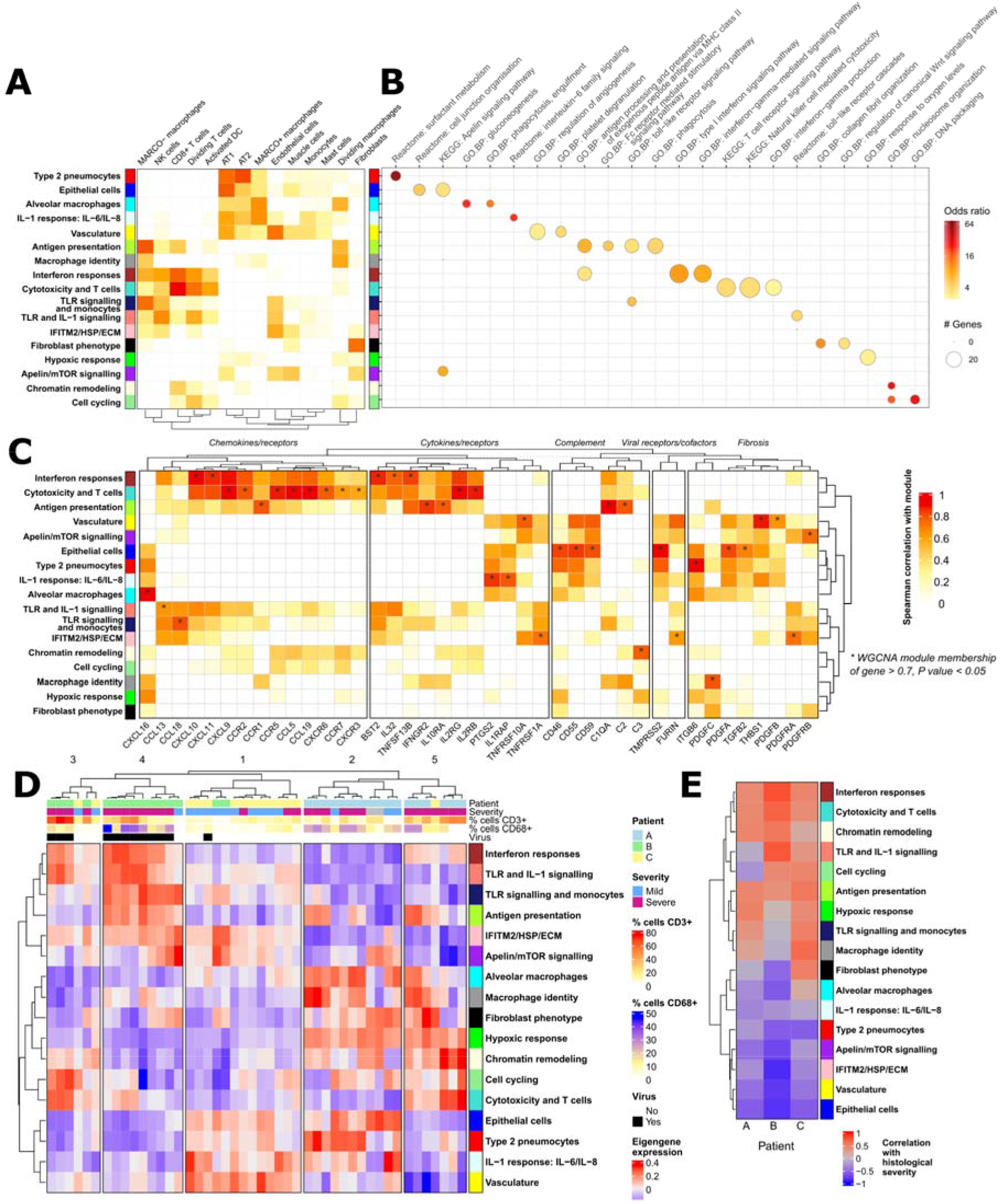
Identification and characterization of gene modules with spatially heterogenous expression in COVID-19 lung tissue. Application of WGCNA to spatial transcriptomic data identified 17 modules of co-expressed genes (see also Supplementary Figure 3). **(A)** Correlation between estimated cell type abundance (as determined by cell deconvolution see **Supplementary Table 6**) and WGCNA module eigengene expression (all AOIs; positive Spearman correlation values are shown). **(B)** Selected GO Biological processes, KEGG and Reactome pathways significantly over-represented in the detected modules (BH adjusted p values <0.05; one-side Fisher’s exact tests; see also **Supplementary Table 5**). **(C)** Genes were selected for impact on pathology: chemokines/receptors; cytokines/receptors; viral receptors and cofactors; complement and fibrosis. Genes were correlated with the eigengene modules (Spearman’s correlation; asterisks indicate high WGCNA membership >0.7 detailed in **Supplementary Table 4**, p value <0.05). **(D)** WGCNA module eigengene expression is shown for each AOI (see also **Supplementary Table 4**). Sampled AOIs are annotated with patient identity, the severity of damage, adjacent virus antigen presence, and the percentage of CD3^+^ and CD68^+^ cells of total nucleated cells. Hierarchical clustering of the 46 AOIs by expression of the WGCNA module eigengenes identified 5 “spatial groups” with distinct patterns of module expression. **(E)** The severity of tissue damage was correlated to module eigengene expression separately for the AOIs from each patient (Pearson correlation).

As expected for lung tissue, we found a set of stromal modules representing (i) “Epithelial cells” (containing *EPCAM*), (ii) “Type 2 pneumocytes” (containing the surfactant encoding genes *SFTPB, SFTPC and SFTPD*), (iii) fibroblasts (“Fibroblast phenotype”, containing *COL1A1, COL3A2, COL5A1* and *THY1*) and (iv) “Vasculature” (containing *CDH5, THBD, ENG*), which all showed corresponding pathway and cell type associations (**Figure 2A-B and Supplementary Figure 3B**). We found three modules that showed a high correlation with the presence of CD68^+^ cells (**Figure 2B-C and Supplementary Figure 3C**). These comprised (i) an “Alveolar macrophage” module that also displayed high expression correlation with the macrophage receptor *MARCO* and enrichment for the “phagocytosis, engulfment” GO biological process, (ii) a “Macrophage identity” module that also showed high expression correlation with the mannose receptor *MRC1* (a marker of alternative “M2” macrophages) and *SIRPA*, and (iii) an “Antigen presentation” module that showed a high correlation with predicted inflammatory monocyte-derived macrophage (monoMac) cell abundance (**Figure 2A, Supplementary Figure 4D**) along with high expression correlation with *SIGLEC1* and *C1QA/B* (**Supplementary Figure 3B**). We also noted a “IFITM2/HSP/ECM” module that contained *MERTK* and *PDGFRA* and a module with pathway enrichments for “Apelin/mTOR signalling”.

Critical COVID-19 is associated with massive lung immune cell infiltration, and in keeping with this we discovered modules of genes with clear associations with lymphocytes and mononuclear phagocytes. The “Cytotoxicity and T cells” module was associated with the GO “Interferon-gamma production” biological process, as well as the KEGG “T cell receptor signalling” and “Natural Killer mediated cytotoxicity” pathways (**Figure 2B**). The expression of this module was positively associated with predicted presence of CD8^+^ cytotoxic T cells, NK cells and activated dendritic cells (DCs) (**Figure 2A**) as well as the presence of CD3^+^ cells (**Supplementary Figure 3C**). This module contained known T cell markers including *CD3D/E, CD2* and *CD8A* as well as genes associated with cytotoxicity such as *PRF1* and *GNLY* (**Supplementary Figure 3B**). It also showed a high correlation with the expression of chemokines including *CXCL9, CXCL10* and *CXCL11*, and had the highest correlation with the expression of *IFNG* (**Figure 2C**) as well as the “Interferon responses” module (**Supplementary Figure 3D**). The “Antigen presentation” module was associated with areas of high CD68 expression by immunofluorescence microscopy (**Supplementary Figure 3C**) and showed positive associations with the “Macrophage identity”, “Cytotoxicity and T cells”, “Interferon response” and TLR signalling related gene modules (**Supplementary Figure 3D**). The “TLR signaling and monocytes” module contained the gene encoding the classical monocyte marker *CD14* along with *CD163* (**Supplementary Figure 3B**). In addition, we discovered several modules that contained genes associated with innate inflammation including an “Interferon responses” module, a “TLR and IL-1 signaling” module and an “IL-1 response: IL-6/IL-8” module that were named according to their pathway enrichments (**Figure 2B**) and correlations with chemokine/cytokine expression (**Figure 2C**). The “Vasculature” and “IL-1 response: IL-6/IL-8” modules contained genes associated with mature neutrophils (*ELANE, MME, MPO, CTSG*) (Supplementary **Figure 3B**). The “Vasculature” module was also associated with the GO.BP platelet degranulation pathway (**Figure 2B)**. Finally, we found three modules that were associated with more general cellular processes. The presence of these modules, which we termed “Cell cycling”, “Chromatin remodelling” and “Hypoxic response” was indicative of immune cell proliferation and oxygen stress.

### Subhead 3: Severe alveolar damage in COVID-19 is linked with myeloid cell antigen presentation, T cell cytotoxicity and expression of the CXCL9/10/11 interferon response genes

We next sought to better understand the variation in cellular and immune phenotype that was present across the tissues and sampled AOIs. To do so we hierarchically clustered the AOIs by expression of the WGCNA module eigengenes. This analysis revealed 5 groups of AOIs with different transcriptional signatures and associations with severity (**Figure 2D and Supplementary Figure 7**). These spatial groups were numbered according to their association with severe damage: spatial group 1 contained the lowest proportion of severe AOIs while spatial group 5 consists of only severe alveolar damage. We performed a ‘cellular phenotype network analysis’ to investigate the correlations between WGCNA eigengene expression (transcriptional phenotype), chemokine and cytokine expression (immune signaling), and the predicted cell type abundances (cell type identity) in each of the 5 spatial groups (**Supplementary Figure 8 B-F**). This analysis provided evidence that severe tissue damage in these COVID-19 patients involved an ensemble of interacting and proliferating immune cells in which myeloid cells such as inflammatory monoMac and DCs were activated by TLR-mediated signaling, expressed IL-1 and interferon alpha, and presented antigen to cytotoxic lymphocytes driving the production of *IFNG* and a specific cassette of chemokines and cytokines. This cassette included high expression of *CXCL9, CXCL10* and *CXCL11*, factors which are known to act via CXCR3 to promote immune cell chemotaxis, extravasation and activation (*15*), as well as IL-32 which is known to stimulate TNFα and IL-6 secretion from macrophages (*16*), and CCL19 which acts via CCR7 to promote DC and central memory T cell migration (*17*).

### Subhead 4: Cytotoxic lymphocytes, IFNγ signaling, myeloid cell activation and TRAIL are associated with severe DAD and are reproducible features of critical COVID-19

Next, we sought to assess the extent to which of the discovered features of severe DAD might represent common features of critical COVID-19. Within AOIs from the each of three patients, we found that severe DAD was consistently associated with the expression of the “Interferon responses”, “Cytotoxicity and T cells”, “Chromatin remodelling”, “Antigen presentation”, and TLR signaling (“TLR and IL-1 signalling” or “TLR signaling and monocytes”) module eigengenes (**Figure 2E**). The “IL-1 response: IL-6/IL-8” module did not show an obvious correlation with DAD severity, while higher expression of gene modules associated with “Epithelial cells” and “Vasculature” were associated with the lowest DAD scores (**Figure 2E**). To investigate commonalities and differences of the tissue pathology of the three patients in more detail, we repeated the ‘cellular phenotype network analysis’ separately for the AOIs of each patient. This analysis demonstrated that a core association between mono-Mac, CD8^+^ T cells, the expression of IFNγ target chemokines *CXCL9, CXCL10* and *CXCL11*, the expression of *IL32*, the expression of the regulator of extrinsic apoptosis *TRAIL (TNFSF10*) and the expression of B cell activating factor, *BAFF (TNFS13B*) was present in the lung tissue from all three patients (**Figure 3A-C**).

**Figure 3:**
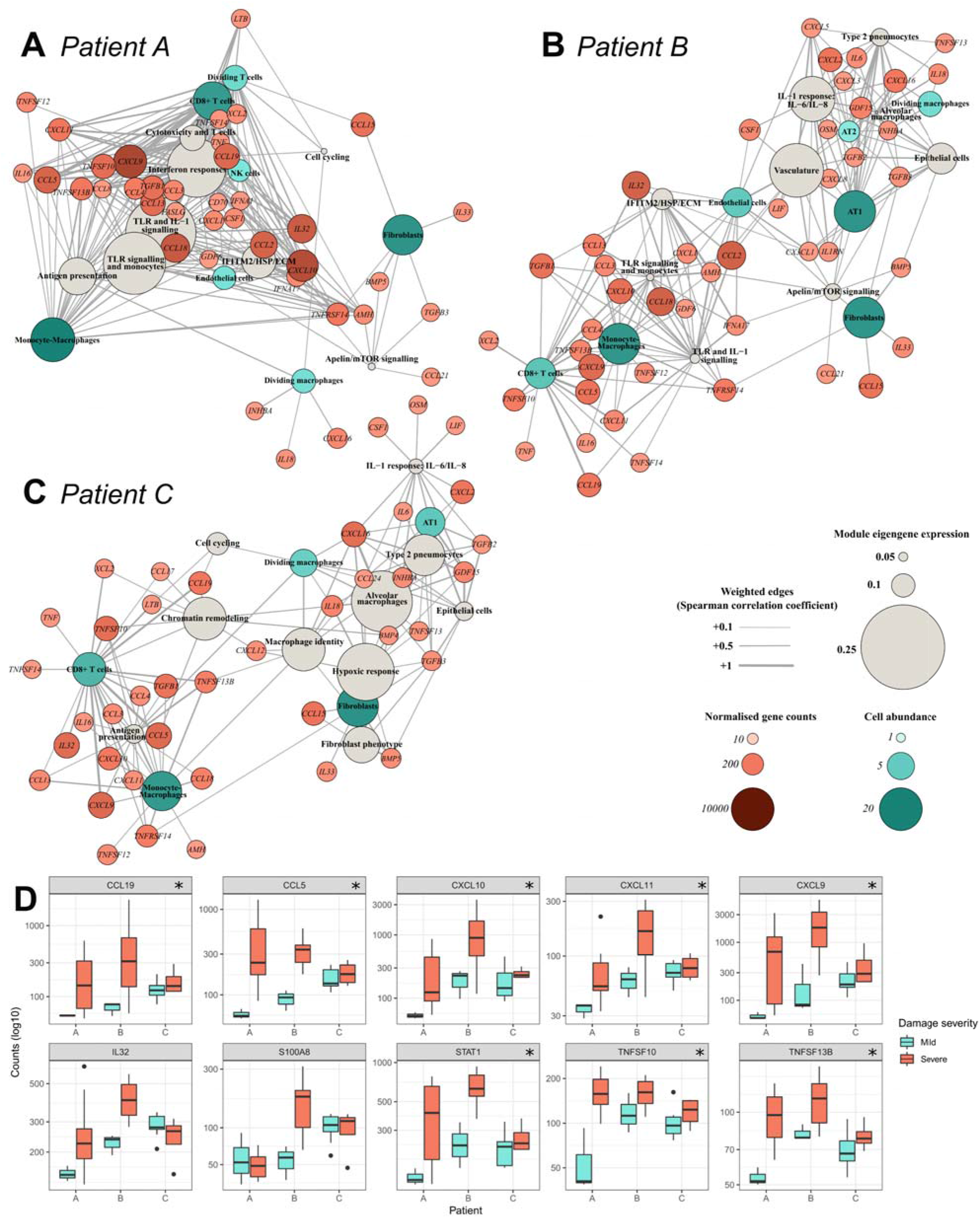
Within patient analysis of spatially associated cellular phenotypes. **(A-C)** The cellular phenotype network analysis diagrams show the correlations (Spearman; p<0.05) between the WGCNA module eigengene expression, predicted cell type abundances and secreted cytokine expression for the three patients (see methods for node inclusion criteria). **(D)** The expression of selected genes in each of the three patients for mild (blue) and severe (red) areas of alveolar damage. Asterisks indicate Benjamini-Hochberg (BH) adjusted *p* value <0.05 in differential expression analysis between mild and severe areas of damage.

To help prioritise immune signalling factors involved in severe DAD we ranked secreted immune signaling factors (and *STAT1*) according to their expression level and up-regulation in the severe areas. In addition to the above signalling molecules, the analysis also highlighted the possible involvement of *STAT1, CCL5, CCL18, CCL13, CCL4, CCL3, CCL21, TGFB1, CXCL12* and *CXCL2* (**Supplementary Figure 9**). To refine our understanding of the cellular sources of involved cytokines we directly correlated their expression with that of specific cell type markers. *CXCL9, CXCL10* and *CXCL11* showed the strongest correlation with CD8^+^ T cell marker genes while *STAT1, TRAIL* (*TNFSF10*) and *BAFF (TNFSF13B*) also showed a strong association with *CD74* and *CD16A (FCGR3A*) which are associated with non-classical monocytes and have been used as markers of inflammatory mono-Mac in COVID-19 (*7*) (**Supplementary Figure 9 and 10)**.

The elevated expression of *TRAIL* in regions of severe DAD is consistent with a previous reports of apoptosis pathway upregulation in alveolar areas in late stage COVID-19 (*6*). The correlation of antigen presentation related genes with DAD severity (**Figure 2E and Supplementary Figure 8**) suggested a directed cytotoxic response, but we did not find an obvious association between the severity and the levels of SARS-CoV-2 N protein RNA (**Supplementary Figure 1E**). This suggested that CD8T/NK cell activation might be triggered by APC presentation or PRR (e.g. TLR) recognition of viral antigens from abortive infection, of self-antigens or of DAMPs or a combination thereof. In support of the possible involvement of DAMPs, we noted that endogenous DAMP encoding genes such as *CCL5* showed robust (if not necessarily up-regulated) expression in areas with severe DAD. Finally, we inspected the expression of key genes of interest with the mild and severe AOIs of each patient. This analysis confirmed a consistent and significant elevation of *CCL19, CCL5, CXCL9, CXCL10, CXCL11, STAT1, TRAIL (TNFSF10), and BAFF (TNFSF13B*) in the areas of severe DAD in the three patients examined (**Figure 3D**). *IL32* showed elevated expression in the severe AOIs of two of the patients while *S100A8* only showed increased severity associated expression in patient B, where its expression was likely neutrophil derived (**Supplementary Figure 6**).

To determine if IFNγ signalling, the discovered *CXCL9/10/11* containing cytokine cassette, cytotoxic lymphocytes, *TRAIL, BAFF* and endogenous DAMP expression are replicable features of lung pathology in critical COVID-19 we investigated the expression of the relevant genes in available community datasets. Examination of single cell data from bronchoalveolar lavage samples (BALF) (*18*) confirmed an increase in *IFNG* expression in T and NK cells in COVID-19 relative to healthy controls along with increased expression of *CXCL9, CXCL10, CXCL11* and *BAFF* in myeloid cells and neutrophils (**Figure 4A**). We also noted a broad increase in *STAT1* and *TRAIL* expression along with induction of *CCL3* and *CCL4* expression in macrophages and neutrophils. The cytotoxic molecule encoding genes *GNLY* and *PRF1* were upregulated in NK and T cells and neutrophils, which were not captured in the healthy controls, constituted an additional source of S100A8/9. The cellular sources and expression levels of these genes were consistent with those observed in other lung tissue single-nuclei and BALF single cell datasets (**Supplementary Figure 11A, B**). Inspection of a previously published spatial analysis of alveolar tissue (which did not discriminate between mild and severe DAD) confirmed up-regulation of *STAT1, CXCL10, BAFF* and *TRAIL* in COVID-19 (**Supplementary Figure 11C**). Increased expression of these genes in COVID-19 was also observed, albeit weakly, in a bulk analysis of macrophages purified from BALF (**Supplementary Figure 11D**). Together, these previously published observations and our targeted spatial analysis suggest the existence of a cellular circuit in which IFNγ production by T/NK cells drives CXCL9/10/11 and BAFF production by myeloid cells in COVID-19. If such a circuit exists, we reasoned that the expression levels of these cytokines should co-vary in the relevant cell types in lung tissue across COVID-19 individuals. To investigate this possibility, we analysed single-nuclei data from the lung tissue of n=16 SARS-CoV-2 infected COVID-19 autopsy donors (*7*) (**Supplementary Figure 12**). In keeping with the existence of the proposed circuit, levels of *IFNG* in per-donor T/NK nuclei pseudobulks showed significant correlations (p <0.05) with the levels of *TRAIL, BAFF and CXCL10* in the paired myeloid nuclei pseudobulks (**Figure 4B**).

**Figure 4:**
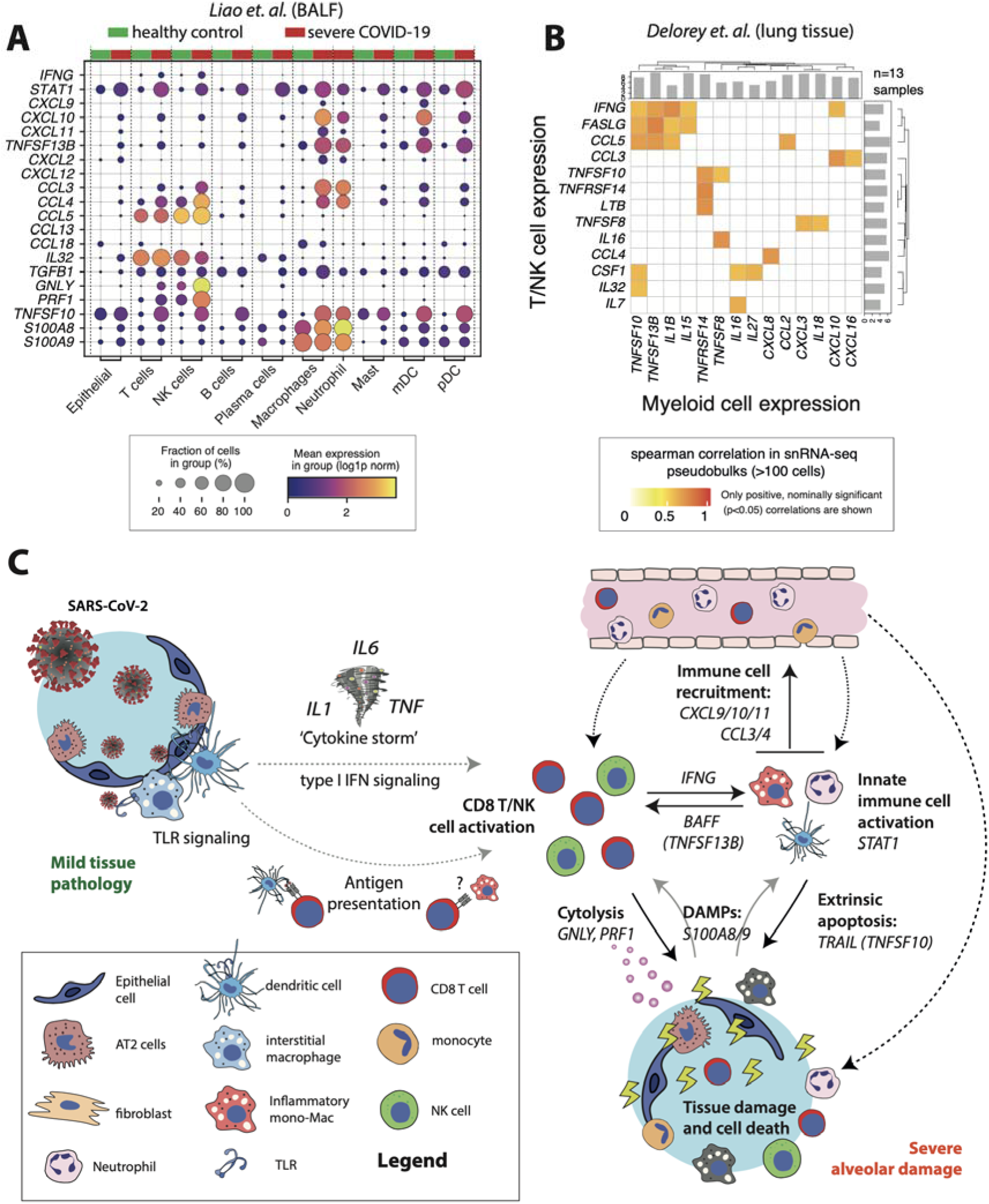
IFNγ production by cytotoxic lymphocytes is associated with severe tissue damage in COVID-19. **(A)** Analysis of the expression of genes of interest in a published bronchoalveolar lavage fluid (BALF) sample single cell dataset comprised of n=4 healthy donors and n=6 severe COVID-19 patients (*18*). **(B)** Correlations between cytokine expression in T/NK and myeloid nuclei pseudobulks constructed from a published single-nuclei atlas of the lung tissue of SARS-CoV-2 infected COVID-19 autopsy donors (*7*). **(C)** Proposed cellular model of severe tissue damage in COVID-19.

## DISCUSSION

Within the lung, COVID-19 manifests in a wide spectrum of DAD and fibroproliferation. The histopathological features are non-specific and there are no clear findings that differentiate SARS-CoV-2 from a number of other respiratory viral infections, particularly those developing following infection with other betacoronaviridae such as SARS-CoV and MERS-CoV (*19–24*). While much has been learnt from analysis of blood and BALF (*10, 18, 25–29*), study of tissue is needed to understand the cellular causes of COVID-19 associated lung damage. Further insight has been gained from single-cell and single nuclei approaches (*7, 30*) but these approaches do not retain spatial information, which is vital for deciphering the interplay between different cell types. Initial applications of spatial proteomics and transcriptomics have started to reveal the spatial landscape of lung damage in COVID-19 (*6, 7*) but molecular details of the signalling circuits that perpetuate pathology remain to be fully elucidated. In this study, we sought to generate new insights by applying network-based analysis approaches to the analysis of rich spatial transcriptomics data generated with Nanostring GeoMx DSP platform. To do so, we used correlation networks to integrate WGCNA module eigengenes, cytokine gene expression levels and computationally predicted cell type abundances. Use of this flexible and extensible ‘cellular phenotype network analysis’ approach uncovered new links between the cell types, biological pathways and cytokines that are associated with lung tissue damage in severe COVID-19.

Our investigations found substantial differences in the cellular and molecular pathology of the lung tissue sampled from the three patients we studied. Tissue from patient A displayed a stronger signature of type 2 pneumocyte and alveolar macrophages. This patient also had the highest expression of a gene module associated with hypoxic response, an observation which was not unexpected given the detection of low oxygen response and p53 stress pathways in the BALF of critical COVID-19 patients (*31*). In contrast with that from the other two cases, the tissue from patient B showed the strongest interferon response and TLR and IL-1 signalling signatures which corresponded with immunohistochemical evidence of high levels of viral infection and the presence of neutrophils. This is consistent with the idea that neutrophil extracellular trap formation (NETosis) may contribute to ongoing inflammation in some COVID-19 patients (*32*). Finally, the samples from patient C showed a lower expression of interferon and hypoxic response signatures and were distinguished by elevated expression of a vasculature associated gene module. Despite these broad differences, our initial within patient analysis revealed a shared association of severe DAD in COVID-19 with interferon signalling, cytotoxicity and T cells, cell proliferation and antigen presentation related genes.

To further elucidate common features of severe COVID-19 we performed within-patient ‘cellular phenotype network analysis’ and explored the reproducibility of our findings using data from published single cell, single nuclei and spatial transcriptomic studies of COVID-19 tissues (*6, 7, 18, 25*). The results provide the basis of a model of severe lung tissue damage in COVID-19 in which IFNγ production by CD8+T and NK cells (i) activates macrophages and other innate immune cells and (ii) induces expression of CXCL9/10/11 (*15, 33*) promoting further recruitment of CXCR3^+^ immune cells (including NK and cytotoxic T cells) in into lymphoid-rich areas (**Figure 4C**). The presence of cytotoxic lymphocytes and elevated expression of *TRAIL* (*TNFSF10*) suggest that severe lung damage in COVID-19 may involve cytolysis and extrinsically regulated apoptosis. In keeping with this model, myeloid cell dysregulation is a hallmark of severe or progressive COVID-19 infection (*34–37*) and there is strong evidence for increased numbers of macrophages in COVID-19 lung tissue (*6, 7, 30*). An increased ratio of CD14^+^HLA-DR^lo^ inflammatory monocytes to tissue-resident alveolar macrophages has also been noted (*18*) and macrophage hyperactivation by persistent IFNγ production has been previously suggested to be a possible mechanism in COVID-19 (*38*). Consistent with the proposed model, despite the well-characterised peripheral blood T and NK cell lymphopenia (*26*), numbers of CD8+ T cells in COVID-19 lung tissue are comparable to those found in healthy individuals, and higher than those found in pneumonia (*6*). NK cells are less abundant than CD8+ T cells in the lung tissue of COVID-19 patients but appear to show an increase in mild disease that is reduced to, or below, healthy levels in severe cases (*6, 7, 18*). Our data extends these observations by showing that, within the lung tissue, cytotoxic CD8+ T cells can localise to interstitial immune cell infiltrates with inflammatory, likely monocyte-derived macrophages and neutrophils within areas of severe COVID-19 associated damage. Based on our transcriptomic analysis we consider that such regions are also likely to contain cytotoxic NK cells but did not find transcriptional or immunohistochemical evidence for CD4 T cell involvement. Our model is likely to be incomplete because important limitations of our work include the number of patients studied, the targeted panel used for the transcriptomic analysis and the limited spatial resolution of the GeoMx DSP platform. Further large-scale studies of COVID-19 lung tissue using higher-resolution proteomic and whole-transcriptome spatial platforms to complement the throughput and targeted sampling that the GeoMx DSP platform affords will be essential for fully deciphering the fine cellular and molecular details of inflammation and severe tissue damage in this disease (*39*).

A key question that arises from the proposed model is the nature of the upstream mechanism(s) by which CD8+ T and NK cells are stimulated to release IFNγ in areas of severe damage. In a similar circuit proposed by Grant et al. based on the analysis of BALF, activation of SARS-CoV-2 reactive T cells in lung alveoli was proposed to be sustained by continued SARS-CoV-2 infection of recruited monocyte derived macrophages (*25*). However, the fact that only one of the three patients studied here showed convincing evidence of viral infection in the tissue suggests that viral antigens may not be the only trigger for cytotoxic lymphocyte activation in severe cases. In support of this hypothesis, a similar observation was made in a study of lung tissue from fatal COVID-19 cases where “virus-independent immunopathology, rather than direct viral cytotoxicity” was proposed to be a primary pathogenic mechanism (*8*). Additionally, it has also been reported that the CD8+ T cell repertoire is more diverse in BALF from severe COVID-19 patients than it is in moderate cases (*18*). Together, these observations suggest that in severe COVID-19, activation of CD8+ T cells in lung tissue may be spatially uncoupled from, and inappropriately sustained after, clearance of viral infection. Candidate mechanisms for this process include exposure to endogenous DAMPs and/or pro-inflammatory cytokines. Possible endogenous DAMPs include the release of S100A8/9 from macrophages (and/or neutrophils) killed by cytotoxic lymphocytes or TRAIL mediated extrinsic apoptosis if not adequately cleared by phagocytosis (*40*). With respect to the possible involvement of inflammatory cytokines, we noted prominent expression of *BAFF (TNFSF13B*) in regions of severe DAD, where, based on inspection of single cell data (*18*), the most likely source was the myeloid cells. It is known that BAFF can promote CD4^+^ T cell IFNγ production and CD8^+^ T cell cytotoxicity in chronic obstructive pulmonary disease (COPD) (*41*), supporting the concept that it may be part of a positive feedback loop that sustains non-specific cytotoxic lymphocyte activation in COVID-19. It is also likely that elevated levels of IL-1, IL-6 and TNF may contribute to dysregulation of CD8 T cells, NK cells and macrophages as part of the so-called ‘cytokine storm’ (*26, 42*).

Currently deployed therapeutics such as dexamethasone and the anti-IL-6 therapy tocilizumab are effective but not curative in all patients (*43*). While the cases that we examined provided some evidence of a role for IL-6 in areas of milder damage, we did not find an obvious link between expression of this cytokine and severe tissue pathology. Our data, suggests that therapeutic targeting of immune cell recruitment via the CXCL9, CXCL10, CXCL11/CXCR3 axis may be a valuable therapeutic strategy for resolution of inflammation in severe COVID-19 where it has become uncoupled from viral clearance. Systemic and lung tissue upregulation of these IFNγ induced cytokines in COVID-19 has been noted in many studies, and this axis has already been proposed a therapeutic target by others for COVID-19 and other serious diseases including cancer (*6, 10, 15, 18, 26, 38, 44*). Less attention has been paid to BAFF (TNFSF13B) which has been shown to be significantly up-regulated in COVID-19 patient plasma, and for which monoclonal blocking antibodies have been developed (*45, 46*). While further study of the role of this interesting cytokine in COVID-19 is required, our data suggest that it may be a valid therapeutic target in severe cases. In two of the three patients studied we also noted a robust up-regulation of *IL-32* in areas of severe damage, and the role of this cytokine, which has important functions in anti-viral responses and can induce expression of pro-inflammatory cytokines may warrant further study in COVID-19 (*47*). Overall, the heterogeneity of the tissue pathology that we observed both within and between the patients studied underscores a need for personalised approaches where the choice of therapy (or therapies) is guided by careful assessment of the stage of disease and viral clearance. For example, while innate immune activation via treatments such as intranasal IFNB1α/β (*48*) are likely to be important for patients unable to clear the virus, our model predicts that they may exacerbate non-specific lymphocyte and myeloid cell activation and tissue damage in virus-free lung tissue and in patients who remain critical after successful viral clearance. As a more general strategy, our findings strongly support the use of combinatorial regimes that pair therapeutics directly targeting the virus such as molnupiravir, remdesivir, ritonavir and recombinant soluble ACE2 (*49, 50*) with those that can temper the dysregulated host immune responses that drive tissue damage.

## MATERIALS AND METHODS

### Design

To delineate the tissue-specific immune pathology in severe COVID-19, we assessed the transcriptomic profile across a spectrum of DAD in well-preserved tissue samples obtained at the point of death from three COVID-19 patients. Medical records of patients were retrospectively reviewed (*51*) and three COVID-19 patients selected for in-depth analysis based on their clinical manifestation of ARDS (**Supplementary Table 1**), typical COVID-19 histology (**Supplementary Table 2**) with a 4-5 score on the Brescia-COVID Respiratory Severity Scale, and a lung-restricted presence of SARS-COV-2 (absence in heart, liver and kidney biopsies). None of the patients were vaccinated against SARS-CoV-2. At least 14 AOIs from each COVID-19 tissue were selected for analysis, spanning on average 0.2mm^2^ (range: 0.05 to 0.33mm^2^) with exclusion of empty space. Areas were selected to represent the spectrum of alveolar injury within each tissue covering regions of a) mild/moderate injury with some conservation of alveolar architecture and b) severe injury with a loss of alveolar structure and significant inflammation. Selected AOIs occupy alveolar and interstitial spaces, except from A_16 and A_17 containing bronchiolar epithelium. The severity grade of each AOI was confirmed post-hoc by 2 pathologists.

### Ethics statement

This study was approved by the ethics committee of the University of Navarra, Spain (Approval 2020.192) and the Medical Sciences Interdivisional Research Ethics Committee of the University of Oxford (Approval R76045/RE001). Tissues were stored at the John Radcliffe Hospital according to Human Tissue Authority regulations (Licence 12433).

### Patients and tissue processing

Post-mortem lung tissues were obtained through open biopsy at the point of death and processed as described(*51*). In brief, tissues were immediately fixed in neutral buffered formalin for <24 hours and then paraffin embedded. Sections (5μm each) were cut for H&E staining, ISH and DSP analysis. Six pathologists reviewed the histology in and agreed on the gross histological characteristics (**Supplementary Table 2**). RNA was extracted from 4-8 × 5μm sections for quantification of SARS-CoV-2 nucleocapsid (N) and envelope protein (E) transcripts (**Supplementary Table 2**).

### In situ hybridization

In-situ hybridization was conducted using the RNAscope2.5 LS Reagent Red Kit, according to manufacturer’s instructions and using the Leica BOND-RXm system. Deparaffinisation and heat-induced epitope retrieval were performed with BOND Epitope Retrieval Solution 2 (ER2, pH 9.0) for 25 minutes at 95°C.□ Hybridization for SARS-CoV-2 RNA was carried out using the RNAscope 2.5 LS Probe V-nCoV2019-S and checked for quality against slides treated with the Positive Control Probe Hs-UBC and Negative Control Probe-DapB.□

### Immunohistochemistry□for SARS-CoV-2 nucleocapsid protein

Deparaffinisation and heat-induced epitope retrieval were performed on the Leica BOND-RXm using BOND Epitope Retrieval Solution 2 (ER2, pH 9.0) for 30 minutes at 95°C.□ Staining was conducted with the Bond Polymer Refine Detection kit, a rabbit anti-SARS-CoV-2 nucleocapsid antibody (Sinobiological; clone: #001; dilution: 1:5000) and counterstained with haematoxylin.□□□ Whole slide image analysis and virus quantification was done using QuPath software (*52*). Lung tissue was distinguished from empty space through ‘Create thresholder’ function on hematoxylin and then virus positive pixels were quantified using ‘Detect positive staining’ function (downsample factor: 10, gaussian sigma: 5μm, hematoxylin threshold: 0.1 OD units, DAB threshold: 0.3 OD units).

### NanoString GeoMx Digital Spatial Profiling

This technique was carried out according to manufacturer’s recommendations for GeoMx-NGS RNA BOND RX slide preparation (manual: MAN-10131-02). Deparaffinization, rehydration, heat-induced epitope retrieval (ER2 for 20 minutes at 100°C) and enzymatic digestion (1μg/ml proteinase K for 15 minutes at 37°C) were carried on the Leica BOND-RX. Tissues were incubated with 10% neutral buffered formalin for 5 minutes and 5 minutes with NBF Stop buffer. The tissue sections hybridized with the oligonucleotide probe mix (Cancer Transcriptome Atlas and COVID-19 spike-in panel) overnight, then blocked and incubated with PanCK-532 (AE1+AE3, Novus), CD3-647 (UMAB54, Origene), CD68-594 (KP1, Santa Cruz) and DNA dye (Syto13-488; Invitrogen) for 1 hour. Tissue sections were then loaded into the GeoMx platform and scanned for immunofluorescent signal.

After selection of areas of interest (AOIs), UV light directed at each AOI released oligonucleotides that were collected and prepared for sequencing. Illumina i5 and i7 dual indexing primers were added during PCR (4 μL of collected oligonucleotide/AOI) to uniquely index each AOI. AMPure XP beads (Beckman Coulter) were used for PCR purification. Library concentration as measured using a Qubit fluorometer (Thermo Fisher Scientific) and quality was assessed using a Bioanalyzer (Agilent). Sequencing was performed on an Illumina NextSeq 2000 and FASTQ files were processed by the NanoString DND pipeline, resulting in count data for each target probe in each AOI.

### Analysis of immunofluorescent images for cell counts

The number of nuclei, CD3^+^ and CD68^+^ cell counts were determined using CellProfiler software. A pipeline was designed to quantify circular objects within RBG files for each AOI(*53*). Global manual intensity thresholds were set for object identification of nuclei and CD3^+^ cells, whilst CD68^+^ cells were identified by adapted intensity thresholds. The efficacy of object identification for each AOI was visually confirmed.

### Quality control and pre-processing of GeoMx transcript expression data

Quality control and initial data exploration was conducted using the GeoMx DSP Analysis Suite. Sequencing quality per AOI was examined and an under-sequenced area (B_04) with zero deduplicated reads was excluded. Expression of each transcript was measured by 5 or more probes; outlier probes (geomean probe in all AOI/geomean of all probes for a transcript = <0.1; or probe failure in the Grubbs outlier test in over 20% AOIs) were excluded and the remaining probes combined to generate a single (post “biological probe QC”) expression value per gene target per AOI.

We evaluated the performance of two normalization strategies, upper quartile and quantile normalization, by investigating their ability to standardize the expression distributions of housekeeping genes, negative control probes as well as the full expression distribution between the AOIs. The expression values for negative control probes that were not reported as global outliers were appended to the gene expression matrix. Based on this assessment (and the results of subsequent PCA analyses) we proceeded with the quantile normalized expression values. We investigated the influence of known technical and biological factors on the variance structure of the dataset by performing PCA of the log2(n+1) transformed quantile-normalized expression values. This analysis revealed that the first component in the data was associated with the aligned read depth statistic. We therefore corrected the quantile normalized expression values for this technical factor using Limma “removeBatchEffect” function. Finally, to distinguish gene expression from background noise, we modelled the expression distribution of the set of negative probes, defining the median of the negative probe expression values + 2*median absolute deviations as a robust detection threshold. In total we retained 1631 genes that passed this detection threshold in at least 2 AOIs. The normalized expression distributions and sample PCA plots obtained following normalization, aligned read depth correction and expression level filtering are shown in **Supplementary Figure 4**.

### Differential gene expression and over-representation analysis

Differential gene expression was calculated for each gene between areas of mild/moderate and severe alveolar damage using linear mixed models (fixed effect: severity, random variable: patient identity). Significance was estimated using Satterthwaite’s degrees of freedom method and *p* values adjusted using Benjamini-Hochberg methods (R libraries: lme4, lmerTest). Over-representation analysis for Gene Ontology Biological Processes (GO.BP) terms was performed on genes with >1.5 fold change and FDR > 0.05 (R library: clusterProfiler(*54*)) with the parameters: pvalueCutoff = 0.05 and qvalueCutoff = 0.10. Redundant and duplicated pathways were removed for **Figure 1D**; for full list of pathways see **Supplementary Data**.

### Weighted gene correlation network analysis (WGCNA)

WGCNA was applied to the log2(n+1) transformed, quantile normalized, aligned read corrected and filtered expression values. The parameter values were set as follows: minimum fraction of non-missing samples for a gene to be considered good=0.5, minimum number of non-missing samples for a gene to be considered good=4, minimum number of good genes=4, cut height for removing outlying samples=100 (no samples removed), minimum number of objects on branch to be considered a cluster=2, network type=signed-hybrid, soft power=4, adjacency correlation function=bicor, adjacency distance function=dist, TOM type=signed, minimum module size (number of genes)=10, dissimilarity threshold used for merging modules: 0.25. The analysis identified 17 distinct modules labelled with colours, plus a module of 3 unassigned genes (grey module). The WGCNA analysis was performed using pipeline_wgcna.py (https://github.com/sansomlab/cornet).

The expression patterns of the modules were summarized by calculation of module eigengenes using the WGCNA package (which defines a modules’ eigengene as the first principal component of the expression of the modules’ gene members). AOIs were hierarchically clustered by expression of the module eigengenes (Pearson correlation distance; optimized leaf ordering) to identify n=5 distinct groups (R “cutree” function). The over-representation of Gene Ontology (GO) categories, KEGG pathways and Reactome pathways in module gene members was tested using one-sided Fishers exact tests (https://github.com/sansomlab/gsfisher) using the union of gene members from all the modules as the background geneset. Representative genesets and pathways that showed significant (BH adjusted p<0.05) over-representations are displayed in the figures. WGCNA modules were further characterized by assessing the correlation of the module eigengenes with a) histological severity (Pearson correlation) b) predicted cell abundances (as estimated using Spatial Decon; Spearman correlation) and c) that of selected genes (Spearman correlation, R library Hmisc).

### Cell deconvolution of GeoMx transcript expression data

Deconvolution of cell types from the gene expression data was performed using the SpatialDecon R library(*14*). For this analysis the full matrix of quantile normalized gene counts (without correction), mean negative probe counts and the cell profile matrix ‘Lung_plus_neut’ were applied as input. The cell profile matrix retrieved from the package was generated from the Human Cell Atlas lung scRNAseq dataset and appended with neutrophil profiles as described in Desai et al. (2020)(*5*). For correlation analyses, the abundance of cell types is reported for cell types present with > 2 relative abundance in > 2 AOIs. Complete cell type output is shown in **Supplementary Figures 5, 7, 9, 10**.

### Construction of AOI group correlation networks

We constructed correlation networks to investigate the relationship between the WGCNA modules, the estimated cell abundances (from SpatialDecon) and the expression levels of immune signalling genes for each of the n=5 AOI spatial groups (R igraph library; layout = layout_with_dh). Nodes were scaled in size and/or colour according to cell abundance, normalized gene expression or WGCNA module eigengene expression. Drawn edges represent significant correlations (p < 0.05) and were weighted according to the value of the correlation coefficient (Spearman’s Rho). WGCNA modules were included in the network if they had a median module eigengene expression >0 within the relevant group of AOIs. Immune signalling genes from the KEGG ‘cytokine-cytokine receptor interaction’ pathway (hsa04060) (human) were included if showed a median expression above the expression detection threshold (as defined in the pre-processing section) in a given AOI group. Cell type abundance estimates were included if they had a median abundance of >2 the given AOI group. For inclusion in the network, nodes (modules, cell types and genes) had higher median values than the thresholds stated above and at least 1 positive correlation to another node type.

### Mass cytometry imaging

5μm thick FFPE lung tissue section slides were stained with metal-conjugated antibodies [Anti-human CD68, CD4, CD8 and CD15 (Fluidigm)] after antigen retrieval. Intercalator-Ir (Fluidigm) was used to stain DNA. Slides were ablated on Fluidigm Hyperion Imaging System using CyTOF7 software (Fluidigm) and visualized using MCD Viewer (Fluidigm). Images were processed for publication using FIJI (*55*) to despeckle and sharpen the images.

## General

Thanks to David Scoville and Andy Nam at NanoString for their assistance with the GeoMx workflow.

## Funding

A.R.C. is funded by the Oxford–Bristol Myers Squibb Fellowship. J.H. is a KRUK Senior Fellow. F.I. is a Wellcome Trust CRCD Fellow. LPH, LD and GO are supported in part by the NIHR Oxford BRC. The investigators acknowledge the philanthropic support of the donors to the University of Oxford’s COVID-19 Research Response Fund. C.E.A, M.V.E, and I.M are supported by Banco Bilbao Vizcaya (BBVA) Foundation for “Ayudas a Equipos de Investigación Científica SARS-CoV-2 y COVID-19”.

## Author contributions

CEDA, MFLA and IM obtained consent, clinical data and the patient samples. IR and CEDA reported on tissue histology. TKT, K-YH and AT developed resources for virus detection. ARC, LC and FI designed and performed the experiments. SNS designed and supervised the computational analysis. SNS and ARC performed the computational analysis.

ARC, FI and SNS interpreted the results and generated the figures. ARC, SNS and FI acquired funding and oversaw the project. PW, RE, LD, GO and LPH designed, performed and acquired funding for imaging mass spectrometry. ARC, JH, PK, IM and FI conceived the original experiments. FI, SNS and ARC wrote the manuscript with input from all of the authors.

## Competing interests

IM discloses grants from Roche, AstraZeneca, Bristol-Myers, Highlight therapeutics and Genmab. IM consults for AstraZeneca, Genmab, Pharmamar, F-STAR, Numab, Pieris, Boston Pharmaceuticals, Bristol-Myers, Highlight therapeutics, Roche, Boehringer Ingelheim and Amunix.

## Data and materials availability

RBG composite images taken of the immunofluorescent stained areas of interest, the NanoString GeoMx DSP raw sequencing, the processed expression data and the metadata are deposited at GEO series GSE186213 (reviewer access token: ovwrkcwgzzupbkn). RBG composite images of the whole immunofluorescent stained tissue sections will be made available on Zenodo. Annotated scripts for the computational analysis are available from https://github.com/sansomlab/covidlung.

## List of supplementary materials

**Supplementary Figure 1:** SARS-CoV-2 RNA and protein expression.

**Supplementary Figure 2:** Normalization and variance of transcriptomic data across areas of interest (AOIs).

**Supplementary Figure 3:** WGCNA modules correlate with spatially divergent features in cell composition, chemokine expression and transcriptional processes.

**Supplementary Figure 4:** Deconvolution of cell lineages across sampled areas in COVID-19 lung tissue.

**Supplementary Figure 5:** Expression of cell type specific marker genes and their correspondence with the predicted cell type abundances.

**Supplementary Figure 6:** Confirmation of Neutrophil presence in a region with severe damage.

**Supplementary Figure 7:** The expression of selected genes that exemplify key differences between the 5 AOI “spatial groups”.

**Supplementary Figure 8:** Network analysis of the AOI spatial groups.

**Supplementary Figure 9:** Prioritisation of cytokines by expression level and association with severe damage.

**Supplementary Figure 10:** Correlation of cell type marker genes with cytokine expression.

**Supplementary Figure 11:** Expression of severe damage associated immune signaling genes in community datasets.

**Supplementary Figure 12:** Analysis of correlation between cytokine expression in different cell types in COVID-19.

**Supplementary Table 1:** Patient characteristics (xlsx file)

**Supplementary Table 2:** Histological characteristics (xlsx file)

**Supplementary Table 3:** Genes differentially expressed between areas of mild and severe tissue damage and associated pathway over-representation analysis (xlsx file)

**Supplementary Table 4:** WGCNA module gene membership and eigengene expression (xlsx file)

**Supplementary Table 5:** WGCNA module pathway analysis (xlsx file)

**Supplementary Table 6:** Predicted cell type abundances (xlsx file)

## References and Notes

1. E. Speranza et al., Single-cell RNA sequencing reveals SARS-CoV-2 infection dynamics in lungs of African green monkeys. Sci Transl Med 13, (2021).

2. D. Wichmann et al., Autopsy Findings and Venous Thromboembolism in Patients With COVID-19: A Prospective Cohort Study. Ann Intern Med 173, 268–277 (2020).

3. G. Grasselli et al., Pathophysiology of COVID-19-associated acute respiratory distress syndrome: a multicentre prospective observational study. Lancet Respir Med 8, 1201–1208 (2020).

4. K. R. Short, E. Kroeze, R. A. M. Fouchier, T. Kuiken, Pathogenesis of influenza-induced acute respiratory distress syndrome. Lancet Infect Dis 14, 57–69 (2014).

5. N. Desai et al., Temporal and spatial heterogeneity of host response to SARS-CoV-2 pulmonary infection. Nat Commun 11, 6319 (2020).

6. A. F. Rendeiro et al., The spatial landscape of lung pathology during COVID-19 progression. Nature, (2021).

7. T. M. Delorey et al., COVID-19 tissue atlases reveal SARS-CoV-2 pathology and cellular targets. Nature, (2021).

8. D. A. Dorward et al., Tissue-Specific Immunopathology in Fatal COVID-19. Am J Respir Crit Care Med 203, 192–201 (2021).

9. A. C. Borczuk et al., COVID-19 pulmonary pathology: a multi-institutional autopsy cohort from Italy and New York City. Mod Pathol 33, 2156–2168 (2020).

10. D. J. Ahern et al., A blood atlas of COVID-19 defines hallmarks of disease severity and specificity. medRxiv, 2021.2005.2011.21256877 (2021).

11. V. Marx, Method of the Year: spatially resolved transcriptomics. Nat Methods 18, 9–14 (2021).

12. C. R. Merritt et al., Multiplex digital spatial profiling of proteins and RNA in fixed tissue. Nat Biotechnol 38, 586–599 (2020).

13. P. Langfelder, S. Horvath, WGCNA: an R package for weighted correlation network analysis. BMC Bioinformatics 9, 559 (2008).

14. P. Danaher et al., Advances in mixed cell deconvolution enable quantification of cell types in spatially-resolved gene expression data. bioRxiv, (2020).

15. R. Tokunaga et al., CXCL9, CXCL10, CXCL11/CXCR3 axis for immune activation - A target for novel cancer therapy. Cancer Treat Rev 63, 40–47 (2018).

16. F. Ribeiro-Dias, R. Saar Gomes, L. L. de Lima Silva, J. C. Dos Santos, L. A. Joosten, Interleukin 32: a novel player in the control of infectious diseases. J Leukoc Biol 101, 39–52 (2017).

17. Y. Yan et al., CCL19 and CCR7 Expression, Signaling Pathways, and Adjuvant Functions in Viral Infection and Prevention. Front Cell Dev Biol 7, 212 (2019).

18. M. Liao et al., Single-cell landscape of bronchoalveolar immune cells in patients with COVID-19. Nat Med 26, 842–844 (2020).

19. Z. Xu et al., Pathological findings of COVID-19 associated with acute respiratory distress syndrome. Lancet Respir Med 8, 420–422 (2020).

20. L. M. Barton, E. J. Duval, E. Stroberg, S. Ghosh, S. Mukhopadhyay, COVID-19 Autopsies, Oklahoma, USA. Am J Clin Pathol 153, 725–733 (2020).

21. T. Menter et al., Postmortem examination of COVID-19 patients reveals diffuse alveolar damage with severe capillary congestion and variegated findings in lungs and other organs suggesting vascular dysfunction. Histopathology 77, 198–209 (2020).

22. T. J. Franks et al., Lung pathology of severe acute respiratory syndrome (SARS): a study of 8 autopsy cases from Singapore. Hum Pathol 34, 743–748 (2003).

23. R. B. Martines et al., Pathology and Pathogenesis of SARS-CoV-2 Associated with Fatal Coronavirus Disease, United States. Emerg Infect Dis 26, 2005–2015 (2020).

24. D. L. Ng et al., Clinicopathologic, Immunohistochemical, and Ultrastructural Findings of a Fatal Case of Middle East Respiratory Syndrome Coronavirus Infection in the United Arab Emirates, April 2014. Am J Pathol 186, 652–658 (2016).

25. R. A. Grant et al., Circuits between infected macrophages and T cells in SARS-CoV-2 pneumonia. Nature 590, 635–641 (2021).

26. M. F. Osuchowski et al., The COVID-19 puzzle: deciphering pathophysiology and phenotypes of a new disease entity. Lancet Respir Med 9, 622–642 (2021).

27. A. Saris et al., Distinct cellular immune profiles in the airways and blood of critically ill patients with COVID-19. Thorax 76, 1010–1019 (2021).

28. J. Schulte-Schrepping et al., Severe COVID-19 Is Marked by a Dysregulated Myeloid Cell Compartment. Cell 182, 1419–1440 e1423 (2020).

29. E. Wauters et al., Discriminating mild from critical COVID-19 by innate and adaptive immune single-cell profiling of bronchoalveolar lavages. Cell Res 31, 272–290 (2021).

30. J. C. Melms et al., A molecular single-cell lung atlas of lethal COVID-19. Nature 595, 114–119 (2021).

31. B. Sposito et al., The interferon landscape along the respiratory tract impacts the severity of COVID-19. Cell 184, 4953–4968 e4916 (2021).

32. B. J. Barnes et al., Targeting potential drivers of COVID-19: Neutrophil extracellular traps. J Exp Med 217, (2020).

33. I. G. House et al., Macrophage-Derived CXCL9 and CXCL10 Are Required for Antitumor Immune Responses Following Immune Checkpoint Blockade. Clin Cancer Res 26, 487–504 (2020).

34. D. Mathew et al., Deep immune profiling of COVID-19 patients reveals patient heterogeneity and distinct immunotypes with implications for therapeutic interventions. bioRxiv, (2020).

35. N. Vabret et al., Immunology of COVID-19: Current State of the Science. Immunity 52, 910–941 (2020).

36. E. J. Giamarellos-Bourboulis et al., Complex Immune Dysregulation in COVID-19 Patients with Severe Respiratory Failure. Cell Host Microbe 27, 992–1000 e1003 (2020).

37. E. R. Mann et al., Longitudinal immune profiling reveals key myeloid signatures associated with COVID-19. Sci Immunol 5, (2020).

38. M. Merad, J. C. Martin, Pathological inflammation in patients with COVID-19: a key role for monocytes and macrophages. Nat Rev Immunol 20, 355–362 (2020).

39. A. Rao, D. Barkley, G. S. Franca, I. Yanai, Exploring tissue architecture using spatial transcriptomics. Nature 596, 211–220 (2021).

40. W. G. Land, Role of DAMPs in respiratory virus-induced acute respiratory distress syndrome-with a preliminary reference to SARS-CoV-2 pneumonia. Genes Immun 22, 141–160 (2021).

41. S. Gao, J. Chen, J. Xie, J. Wang, The effects of BAFF on T lymphocytes in chronic obstructive pulmonary disease. Respir Res 21, 66 (2020).

42. H. Hachem et al., Rapid and sustained decline in CXCL-10 (IP-10) annotates clinical outcomes following TNF-alpha antagonist therapy in hospitalized patients with severe and critical COVID-19 respiratory failure. medRxiv, (2021).

43. R. C. Group, Tocilizumab in patients admitted to hospital with COVID-19 (RECOVERY): a randomised, controlled, open-label, platform trial. Lancet 397, 1637–1645 (2021).

44. M. Blot et al., CXCL10 could drive longer duration of mechanical ventilation during COVID-19 ARDS. Crit Care 24, 632 (2020).

45. C. Schultheiss et al., Next-Generation Sequencing of T and B Cell Receptor Repertoires from COVID-19 Patients Showed Signatures Associated with Severity of Disease. Immunity 53, 442–455 e444 (2020).

46. Y. Tong, S. Zhong, Z. Shan, W. Yao, H. Tian, A novel human anti-BAFF neutralizing monoclonal antibody derived from in vitro immunization. Biomed Pharmacother 119, 109430 (2019).

47. Y. Zhou, Y. Zhu, Important Role of the IL-32 Inflammatory Network in the Host Response against Viral Infection. Viruses 7, 3116–3129 (2015).

48. E. Sallard, F. X. Lescure, Y. Yazdanpanah, F. Mentre, N. Peiffer-Smadja, Type 1 interferons as a potential treatment against COVID-19. Antiviral Res 178, 104791 (2020).

49. W. Fischer et al., Molnupiravir, an Oral Antiviral Treatment for COVID-19. medRxiv, (2021).

50. A. Zoufaly et al., Human recombinant soluble ACE2 in severe COVID-19. Lancet Respir Med 8, 1154–1158 (2020).

51. B. Recalde-Zamacona et al., Histopathological findings in fatal COVID-19 severe acute respiratory syndrome: preliminary experience from a series of 10 Spanish patients. Thorax 75, 1116–1118 (2020).

52. P. Bankhead et al., QuPath: Open source software for digital pathology image analysis. Sci Rep 7, 16878 (2017).

53. M. R. Lamprecht, D. M. Sabatini, A. E. Carpenter, CellProfiler: free, versatile software for automated biological image analysis. Biotechniques 42, 71–75 (2007).

54. G. Yu, L. G. Wang, Y. Han, Q. Y. He, clusterProfiler: an R package for comparing biological themes among gene clusters. OMICS 16, 284–287 (2012).

55. J. Schindelin et al., Fiji: an open-source platform for biological-image analysis. Nat Methods 9, 676–682 (2012).

