## Supplementary Figures for "Spatial transcriptomic characterization of COVID-19 pneumonitis identifies immune circuits related to tissue injury"

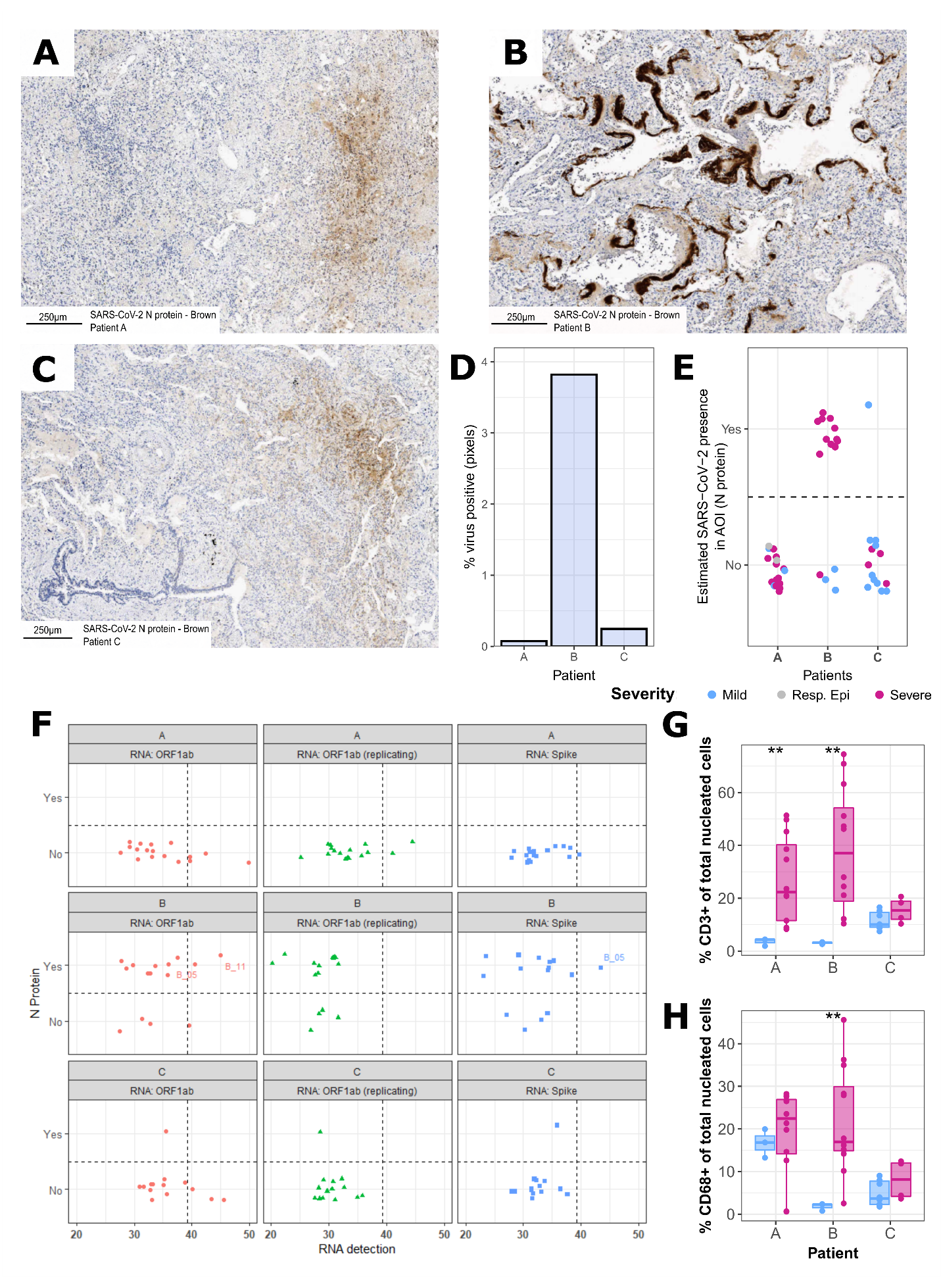

**Supplementary Figure 1: SARS-CoV-2 protein expression.** (**A**-**C**) Examples of immunohistochemistry for the detection of SARS-CoV-2 nucleocapsid protein (brown) in lung tissues from each patient (A-C). Scale bars: 250μm. (**D**) Quantification of the N protein across lung samples (% positive pixels of total tissue pixels). (**E**) Quantification of N protein was conducted on a non-sequential lung section. After alignment of the GeoMx immunofluorescent microscopy and the immunohistochemistry, the presence of SARS-CoV-2 virus was estimated in each AOI and annotated by the severity of alveolar injury (grey circles indicate AOI containing respiratory epithelium). (**F**) Probes against viral SARS-CoV-2 genes (ORF1ab, anti-sense ORF1ab/ORF1abREV and S/Spike) were included in the GeoMx probe panel. Normalised expression values are shown against the presence of SARS-CoV-2 N protein for each patient. The vertical dashed line is the threshold of detection (the median of the negative probe expression values + 2*median absolute deviations). The percentage of CD3^+^ (**G**) and CD68^+^ (**H**) cells of total nucleated cells were derived from the immunofluorescent imaging from mild and severe AOI (mild/moderate-blue, severe-magenta) were compared for each patient (Mann-Whitney test, ** p<0.01).

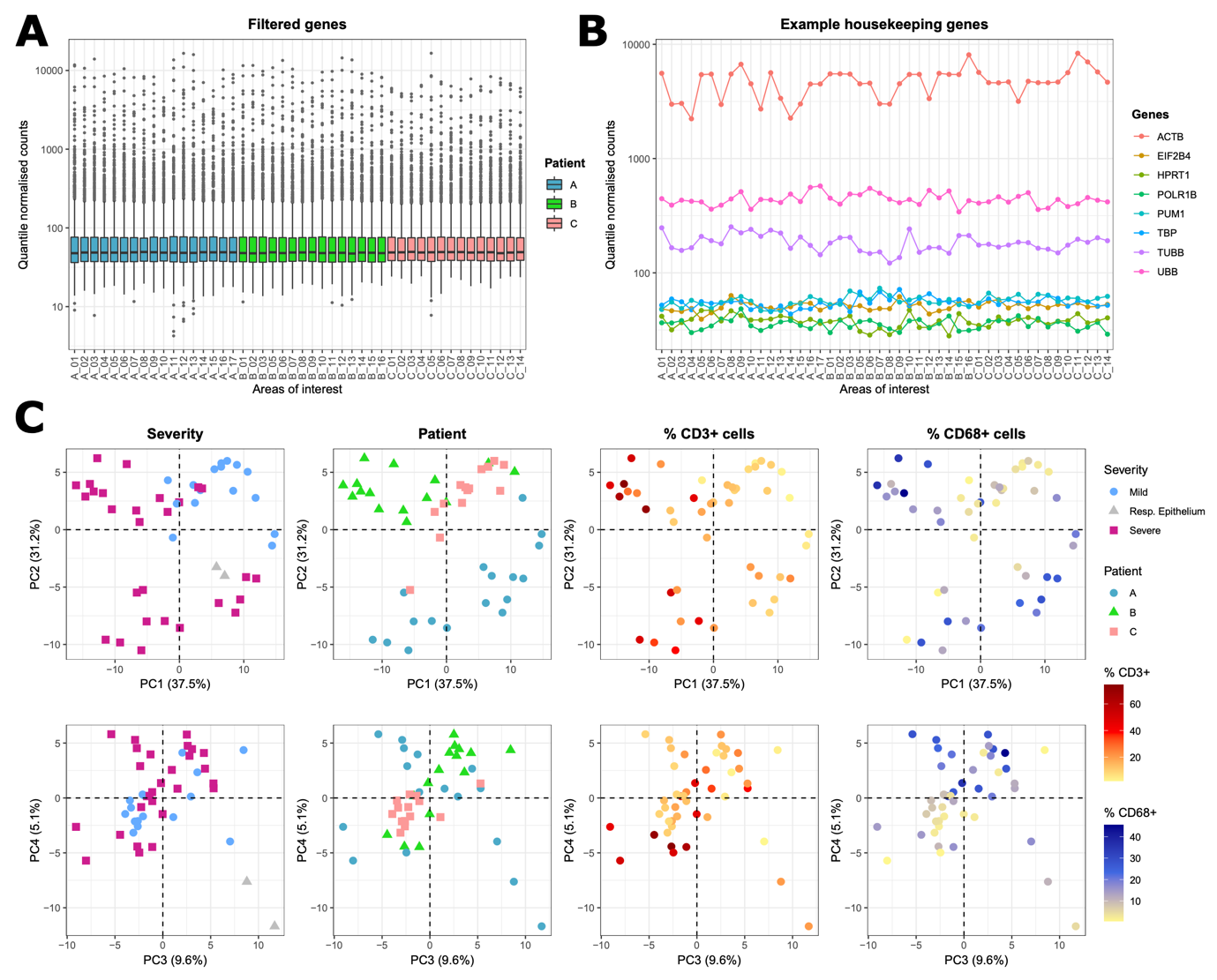

**Supplementary Figure 2: Normalization and variance of transcriptomic data across areas of interest (AOIs).** (**A**) The distribution of all gene and (**B**) selected housekeeping gene expression levels after the application of quantile normalization across the AOIs. (**C**) Principal component analysis of the variance between the sampled AOIs (PCA performed with the pre-processed quantile normalised, log2 transformed, aligned read corrected and expression-level filtered gene expression values). Here PCs 1-4 are shown, annotated by the severity of alveolar damage (severe: pink/square, mild/moderate: light blue/circle, respiratory epithelium: grey triangle), patient identity (A: blue/circle, B: green/triangle, C: pink/square), percentage of CD3^+^ (red gradient) and CD68^+^ cells (blue gradient) of total nucleated cells.

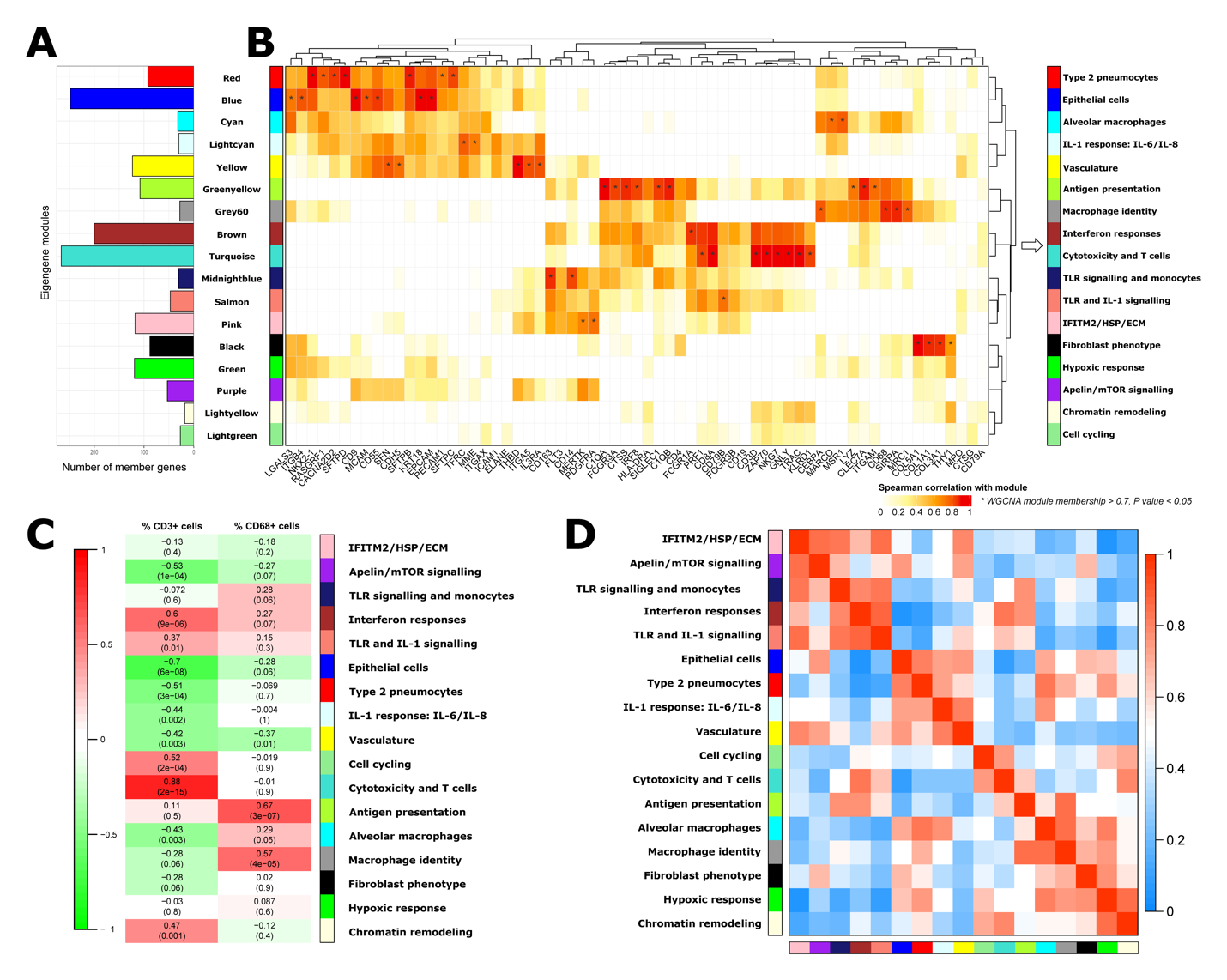

**Supplementary Figure 3: WGCNA modules correlate with spatially divergent features in cell composition, chemokine expression and transcriptional processes.** (**A**) The number of genes present in each module. (**B**) Initial characterisation by exploration of the correlations of cell type marker and immune function genes with the WGCNA module eigengenes (only positive Spearman correlation values are shown; asterisks indicate high module membership > 0.7 detailed in **Supplementary Table 4**, *p* value < 0.05). Simplified module aliases (right) describe each module based on their association with cell lineages, function-associated genes and defining biological pathways. (**C**) Module-trait relationships between the WGCNA gene module eigengenes and the percentage of CD3^+^ or CD68^+^ cells in sampled AOIs. The pearson correlation is shown with the Student asymptotic *p* values given in brackets. (**D**) The heatmaps shows the WGCNA eigengene adjacency.

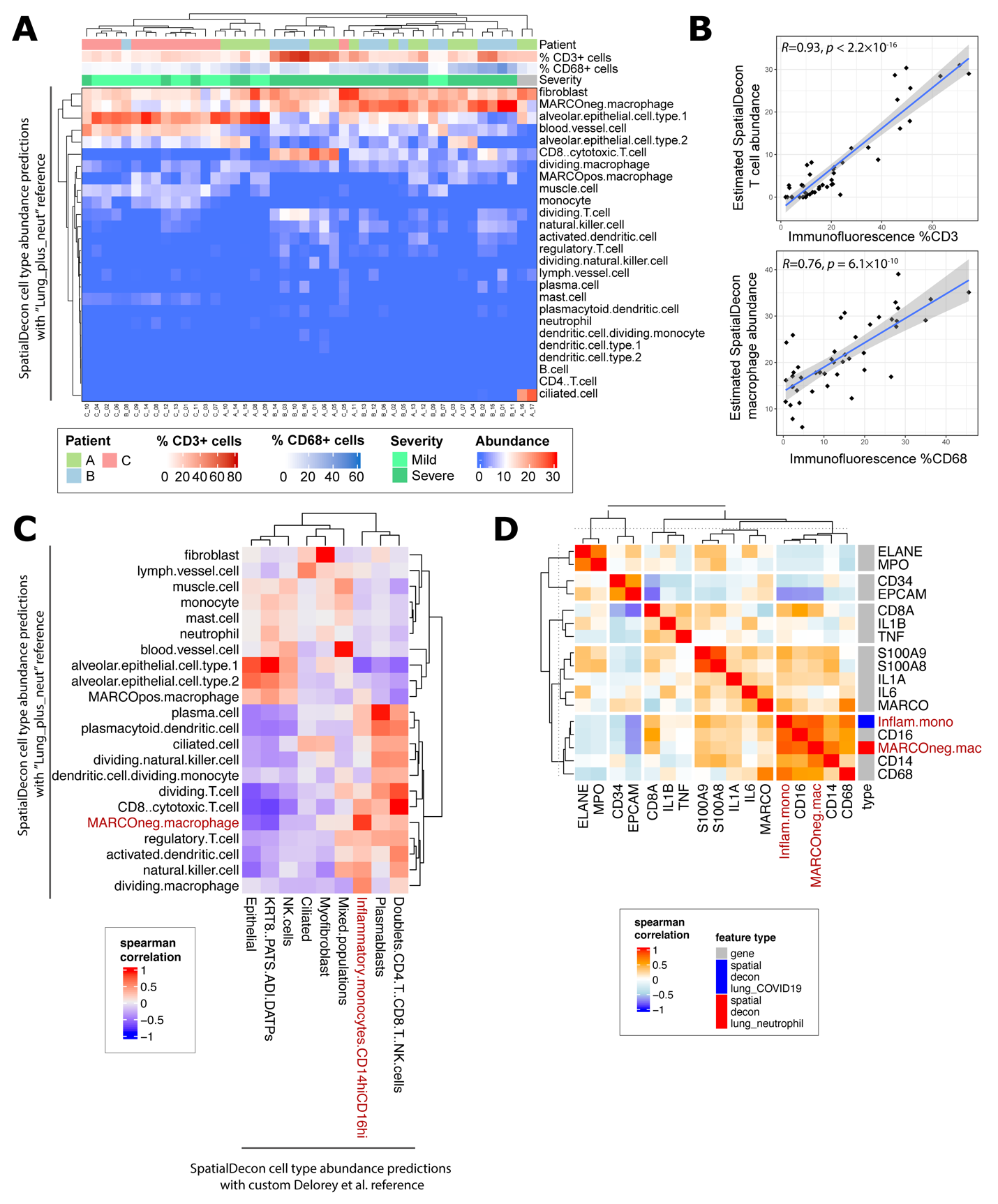

**Supplementary Figure 4: Deconvolution of cell lineages across sampled areas in COVID-19 lung tissue.** (**A**) The abundance of cell types was estimated from the RNA expression data obtained from each sampled area (AOI) across the post-mortem COVID-19 lung tissues (R package SpatialDecon using cell profile matrix derived of healthy lung scRNAseq appended with neutrophil populations ‘Lung_plus_neut’; see also **Supplementary Table 6**). The abundance is hierarchically clustered and annotated with patient identity, the percentage of CD3 or CD68 cells of total nuclei, and the alveolar damage severity. (**B**) The estimated cell abundance is plotted with the percentage of CD3^+^ (top panel) or CD68^+^ (bottom panel) cells of total nuclei, calculated from analysis of the immunofluorescent images (linear model; confidence interval = 0.95). (**C**) The cell type abundances estimated by SpatialDecon decomposition with the ‘Lung_plus_neut’ reference are compared with those from a SpatialDecon decomposition with a custom reference built from the single cell/nuclei analysis of COVID-19 lung tissue in reference (*1*). We noted that the predicted frequencies of “MARCOneg macrophages” from decomposition with the healthy “Lung_plus_neut” reference corresponded closely with those for “Inflammatory monocytes CD14hi CD16hi” from the decomposition with the published COVID-19 reference data (*1*). (**D**) Correlation of key monocyte and macrophage marker genes with the predicted “MARCOneg macrophages” (“Lung_plus_neut” reference) and ”Inflammatory monocytes CD14hi CD16hi” (*1*) abundances. Based on this analysis we re-named the “MARCOneg macrophages” abundance predictions as “mono-Mac”.

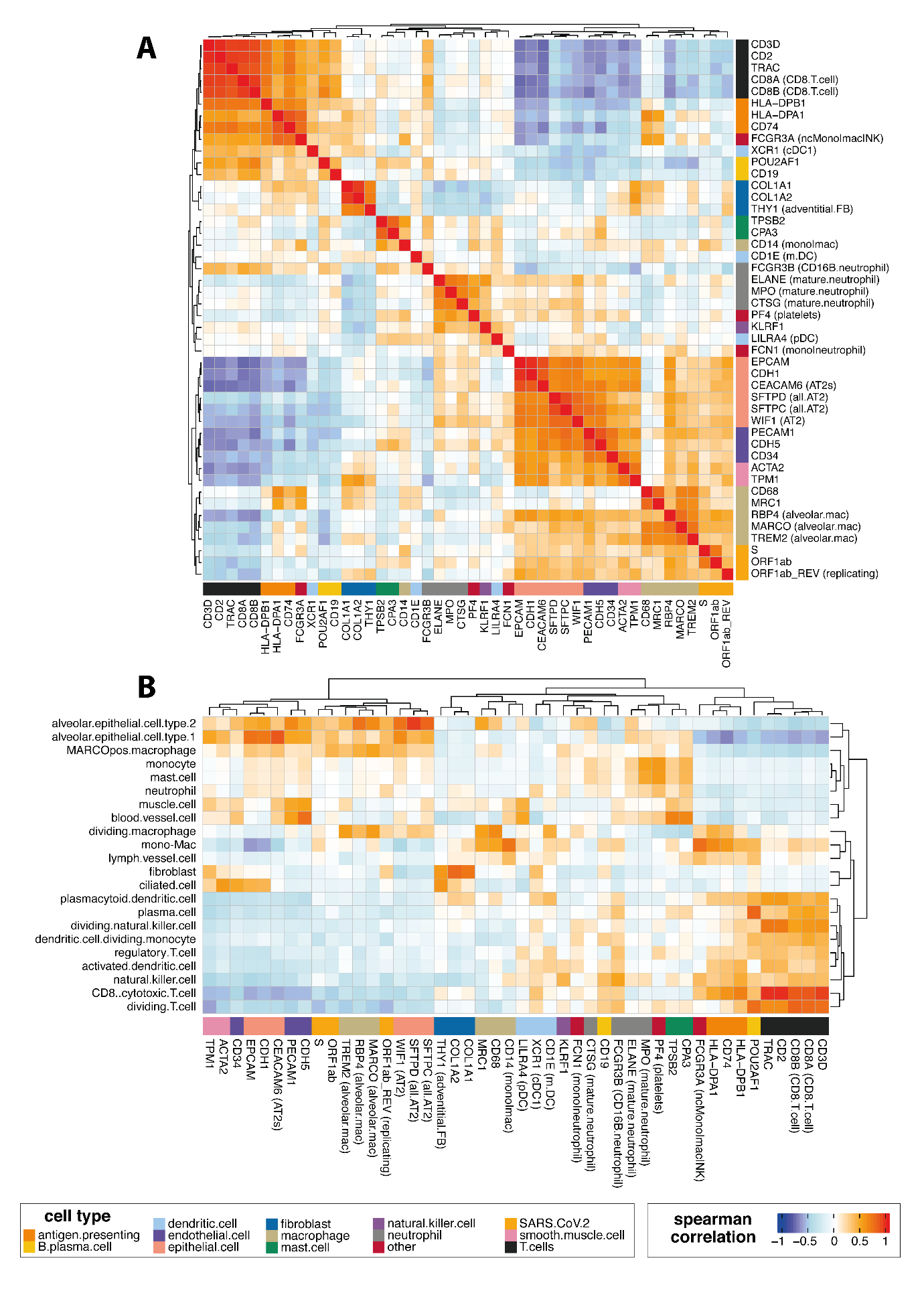

**Supplementary Figure 5: Expression of cell type specific marker genes and their correspondence with the predicted cell type abundances.** (A) Highly specific cell type marker genes were selected from the literature (including from (*2-6*)) and their expression correlated across all the AOIs. The probes designed to capture the viral transcripts are also included. (B) Correspondence between the cell type marker genes and the SpatialDecon predictions.

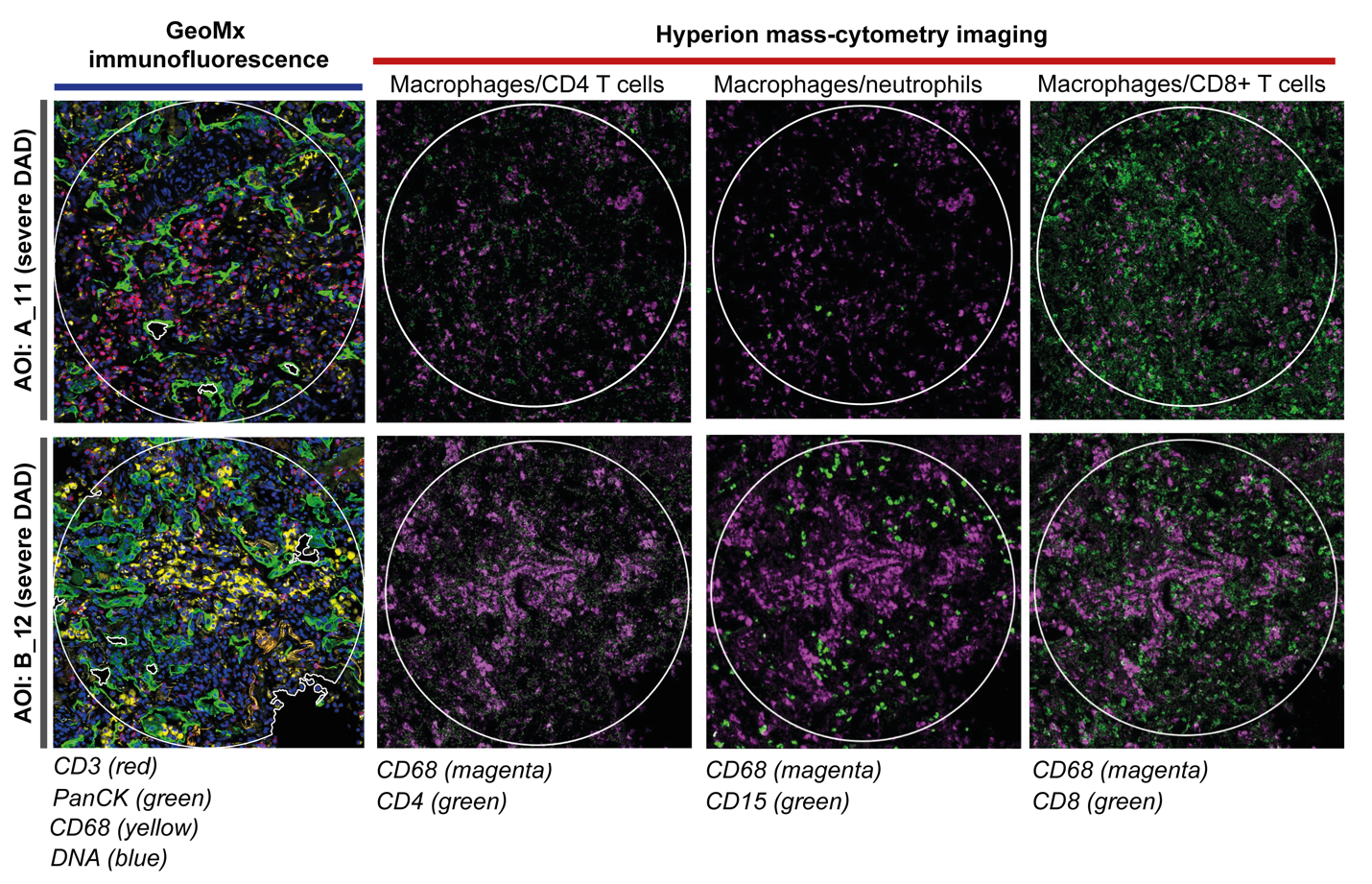

**Supplementary Figure 6: Confirmation of Neutrophil presence in a region with severe damage.** Hyperion imaging of n=2 selected AOIs for expression of CD4 (CD4 T cells), CD8 (CD8 T cells), CD68 (macrophages) and CD15 (neutrophils). Left: IHC images, right: Hyperion mass-tagged antibody images.

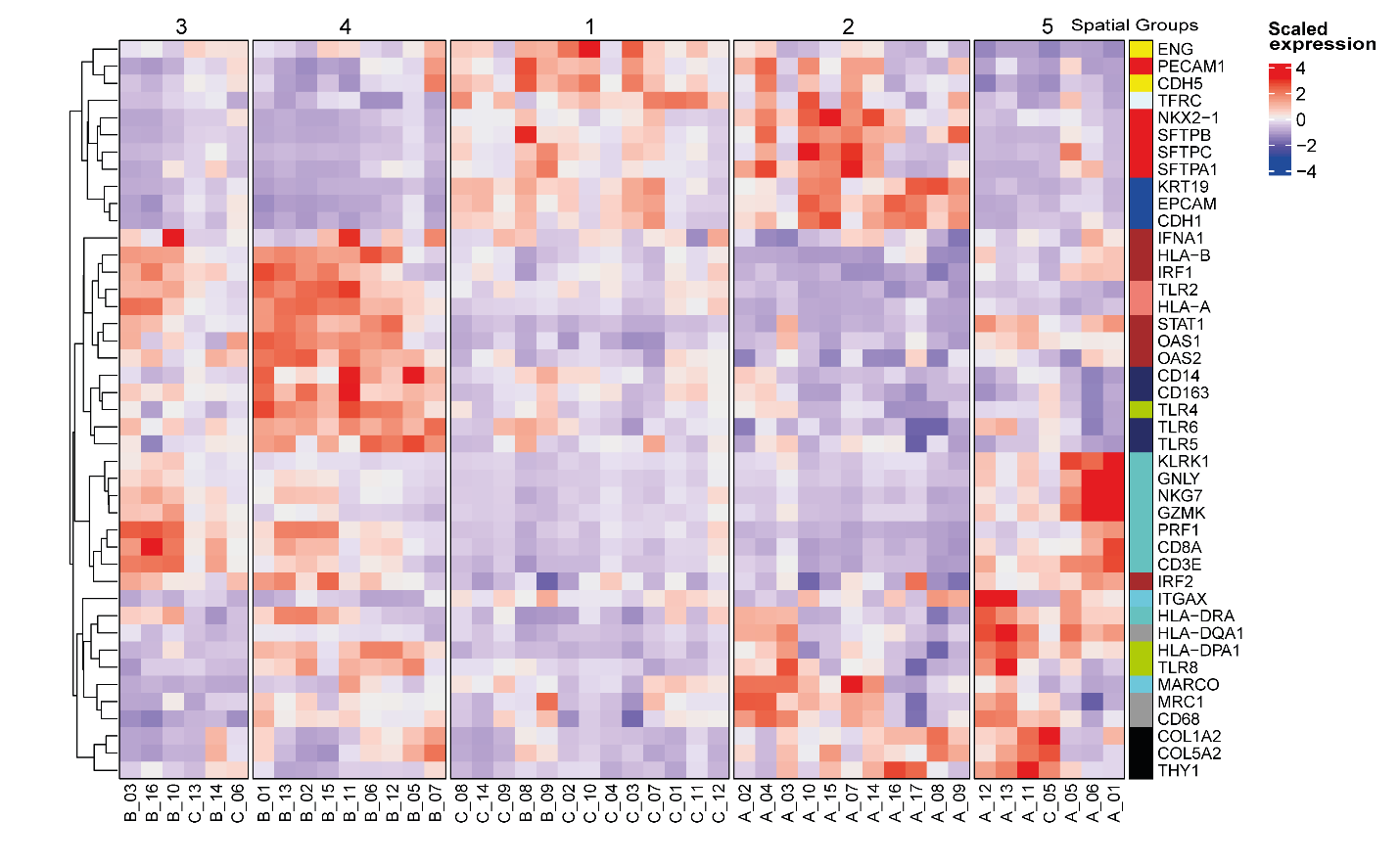

**Supplementary Figure 7:** **The expression of selected genes that exemplify key differences between the 5 AOI “spatial groups”.** See also **Figure 2D**. Spatial group 1 included AOIs from patients B and C that were marked with high expression of the stromal eigengenes (with expression of the vascular genes *ENG, PECAM1, CDH5*; the surfactant genes *SFTPA1, STFPB*, and *SFTBC* and the epithelial genes *KRT19, EPCAM* and *CDH1*), as well as the “IL-1 response: IL-6/IL-8” eigengene, and that were mostly of mild severity. The AOIs in spatial group 2 all came from patient A, showed expression of macrophage and “Hypoxic response” eigengenes in addition to that of many of the stromal signatures in spatial group 1, but also a number of macrophage and antigen-presenting genes (*HLA-DR* genes, *MARCO, MRC1* and *CD68*), and were associated with both mild and severe DAD (**Figure 2D**). These three spatial groups (3, 4 and 5) lacked enrichment of the epithelial eigengenes, shared higher expression of the “Interferon responses” module and were predominantly composed of AOIs with severe DAD and higher fractions of CD3^+^ T cells. AOIs from spatial groups 3 and 5 showed higher expression of the “Cytotoxicity and T cells” and “cell cycling” eigengenes, while the AOIs from spatial group 4 (which were all from patient B) showed higher expression of the “TLR signalling and monocytes” eigengene. Spatial groups 3 and 5 shared expression of NK, CD8, and cytotoxicity genes (e.g. *NKG7, CD8A, GNLY, GZMK*, and *PRF1*), but differed in their expression of interferon genes (spatial group 3) vs. macrophage genes (spatial group 5) (**Figure 2D**).

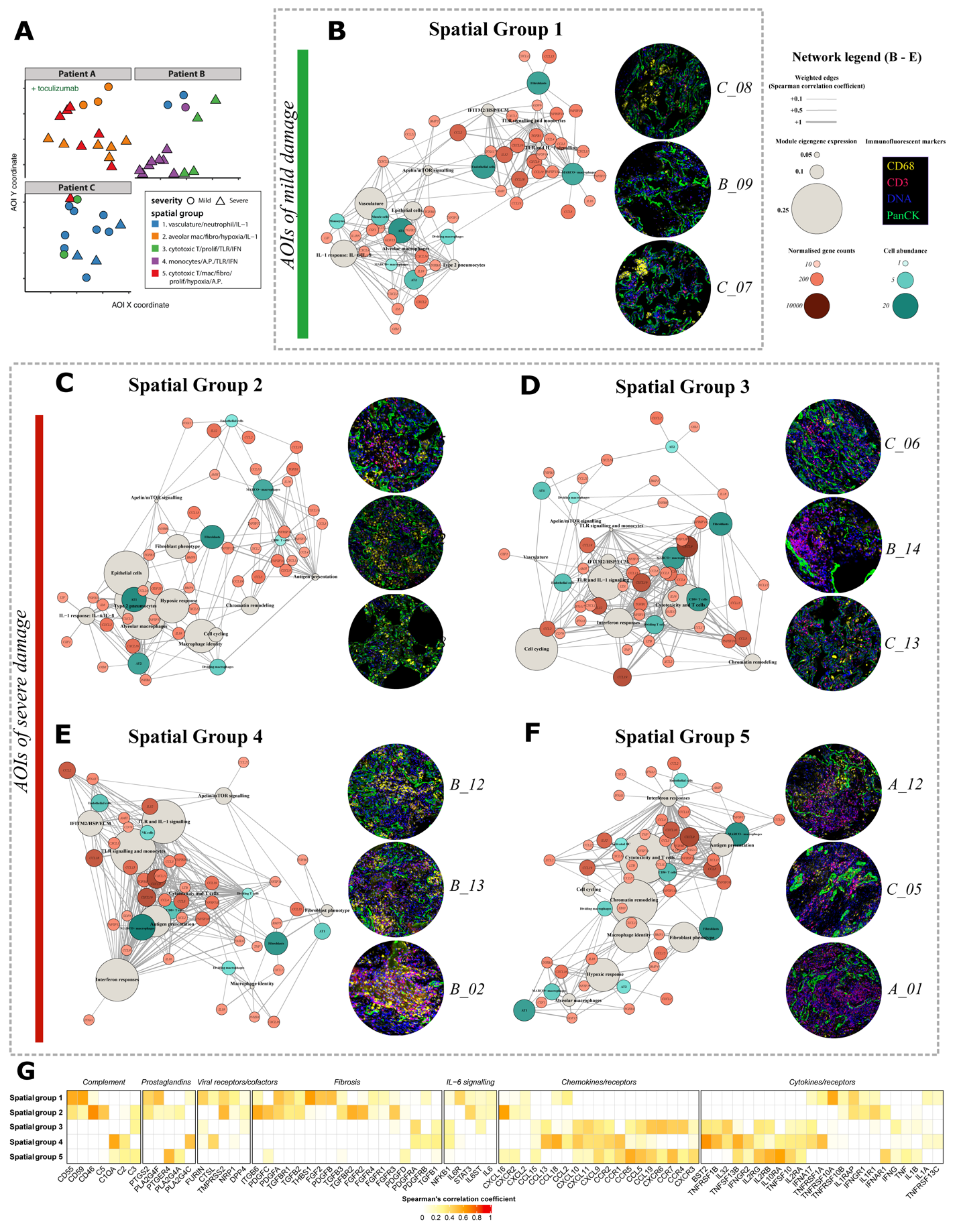

**Supplementary Figure 8: Network analysis of the AOI spatial groups.** (**A**) Locations of the AOIs is shown by patient and AOI group. (**B**) Correlation network for spatial group 1. (**C**) Correlation network for spatial group 2. (**D-F**) Correlation networks for spatial groups 3-5 show associations between (i) expression of the WGCNA modules “Cytotoxicity and T cells”, “Antigen presentation”, “Interferon response”, “Cell cycling” and “Chromatin remodelling”, (ii) predicted abundances of “MARCO- macrophages”, “Activated DC”, “NK”, “CD8 T cells”, “Dividing T cells”, “Dividing macrophages” and (iii) expression of *CCL2/3/4/5/13/18/19*, *CXCL9/10/11, FASLG, IFNA1, IFNA17, IL16/18/32, LTB, TNF, TNFSF10, TNFSF13B and XCL2*. Additinaly the “TLR and IL-1 signalling” module eigengene was implicated in the spatial group 3 and 4 networks from patients B and C while the “IL-1 response: IL-6/IL-8” module was observed in the spatial group 1 and 2 networks from patients A, B and C suggesting that IL-1 signaling is involved in the COVID-19-induced lung tissue damage. In agreement with the computational predictions, immunofluorescent confocal microscopy analysis (representative inset images B-F) confirmed the presence of a large number of T cells in the AOIs with the highest levels of tissue damage. Networks were constructed with selected WGCNA eigengene modules (grey nodes), cytokines and chemokines red gradient nodes) and cell abundance estimates from cell deconvolution (blue gradient nodes) (see methods for node inclusion criteria). Nodes are scaled in size and/or colour to the relative module eigengene expression (grey nodes), the normalized gene counts (red gradient nodes) and the cell abundance (blue gradient nodes). Edges represent positive correlations (Spearman; *p*<0.05) between a gene/cell type and eigengene modules across the whole dataset and are weighted to the Spearman Rho coefficient (+0.1 to +1). Three immunofluorescent images of representative AOI from each spatial group are shown to illustrate major morphological features and cell composition (pan cytokeratin-green, DNA-blue, CD3-red, CD68-yellow). (**G**) Positive correlations between spatial groups and the expression of genes selected for impact on pathology is shown: *chemokines/receptors, cytokines/receptors*, *viral receptors and cofactors*, *IL-6 signalling, complement, fibrosis* and *prostaglandins* (Spearman’s rank correlation, white 0 to red +1).

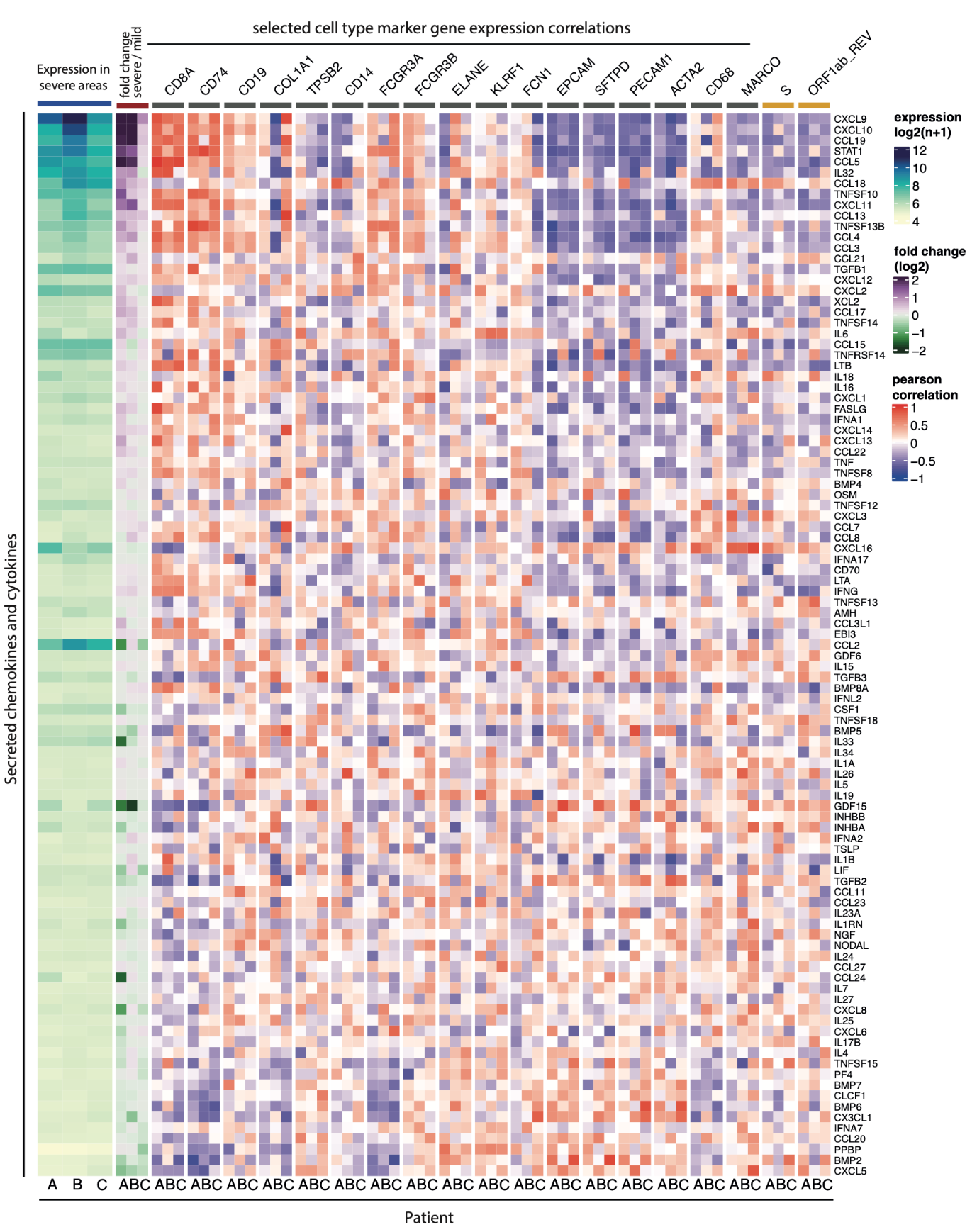

**Supplementary Figure 9: Prioritisation of cytokines by expression level and association with severe damage.** The heat map shows, from left to right (i) the expression level of the cytokines in the three patients in the severe AOIs, (ii) the fold change of the cytokines between the mild and severe areas within each patient, and (iii) the correlations of the cytokines with representative cell type marker genes within each of the patients. The cytokines (rows) are ordered by the sum of the expression rank and fold change rank (such that cytokines with the highest expression and association with severe damage are located at the top of the plot).

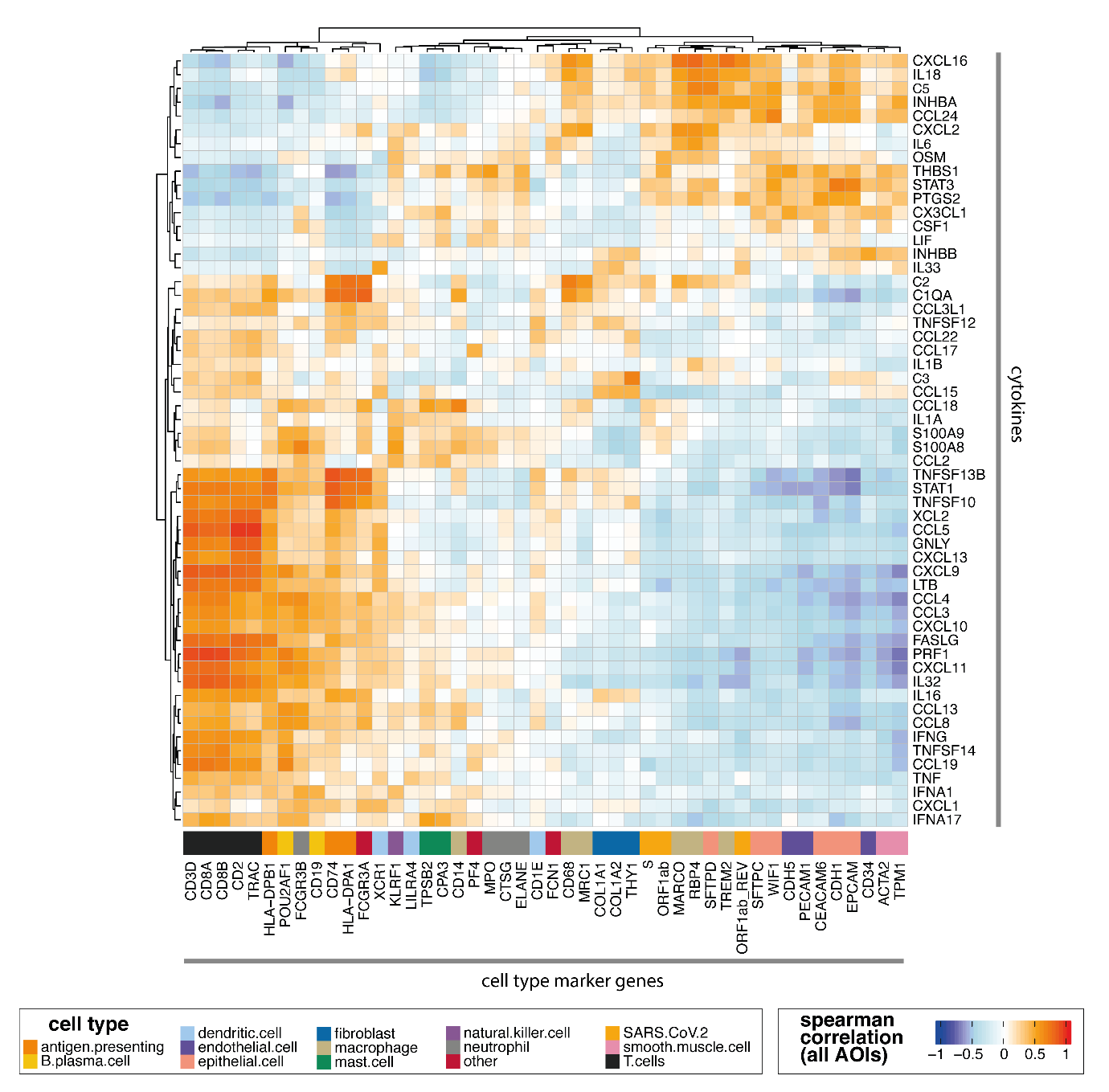

**Supplementary Figure 10: Correlation of cell type marker genes with cytokine expression.** The expression of the cell type marker genes (Supplementary Figure 5) was correlated with the expression of cytokines across all the AOIs.

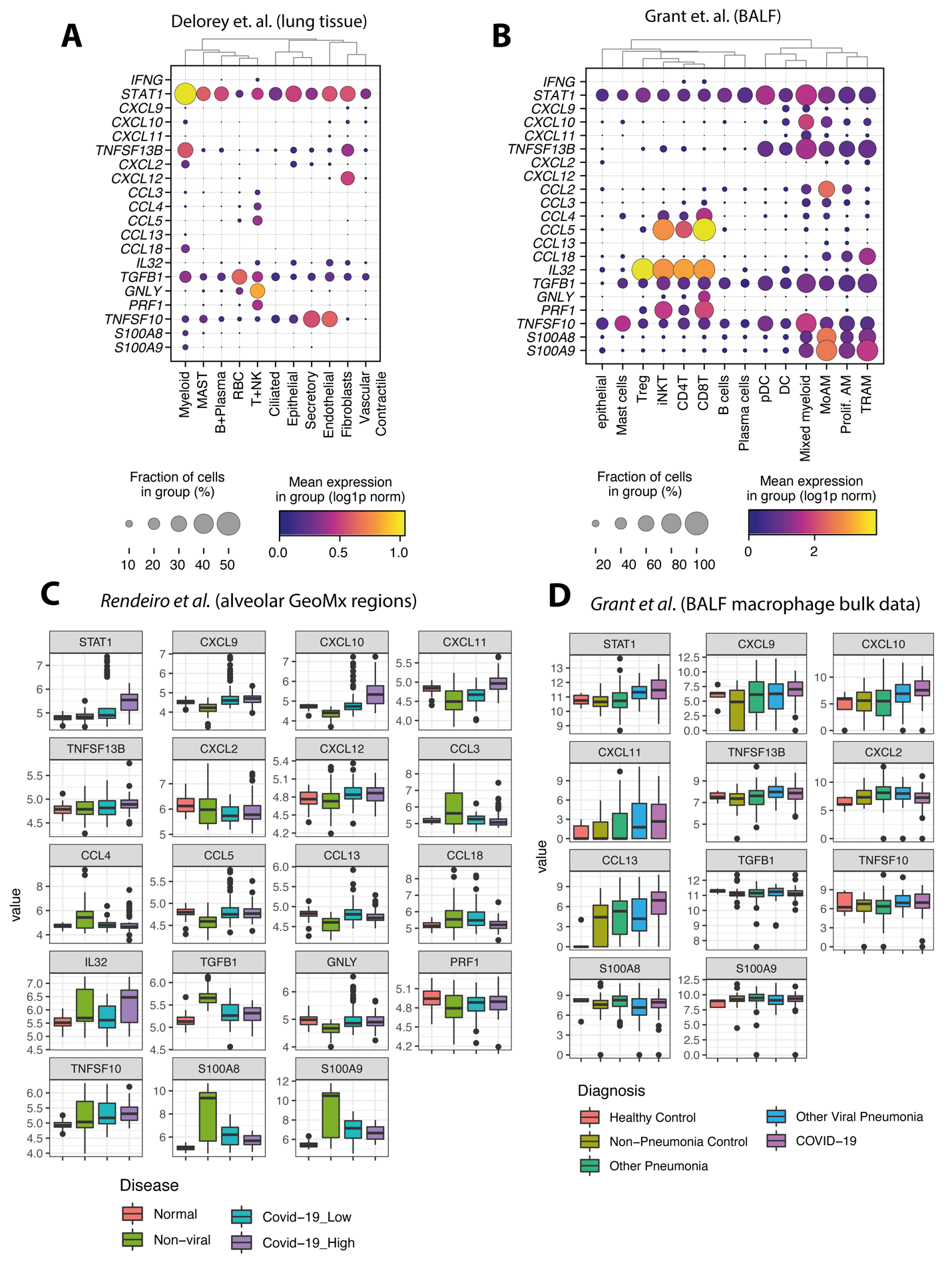

**Supplementary Figure 11: Expression of severe damage associated immune signaling genes in community datasets.** (A) The dotplot shows the expression of the selected genes in the single nuclei cell/data from Delorey et. al. (n=23 lung tissue samples from n=16 SARS-CoV-2 infected COVID-19 autopsy donors; “lung.h5ad.gz” data object retrieved from <https://singlecell.broadinstitute.org/single_cell/study/SCP1052/covid-19-lung-autopsy-samples>) (*1*). Counts were total count normalised and log1p transformed with Scanpy prior to visualisation. Cells in the annotated “Mesothelial”, doublet and “Mixed population” SubClusters were excluded from the analysis. The remaining cells were grouped by the Authors “Cluster” annotations. (B) The dotplot shows expression of the selected genes in the single cell data from (*3*) (n=10 brochoalveolar lavage (BAL) fluid samples from n=10 COVID-19 positive patients). The “GSE155249_main.h5ad.gz” data object was retrieved from <https://www.ncbi.nlm.nih.gov/geo/query/acc.cgi?acc=GSE155249>. For plotting, the original epithelial, CD4 T cell, CD8 T cell, DC, MoAM and TRAM “cell_type” annotations were merged into single categories. (C) The boxplots show the expression of the selected genes in the alveolar tissue GeoMx samples from (*7*). These comprised of n=73 AOI from n=4 “Covid-19_High” patients, n=74 AOI from n=4 “Covid-19_Low” patients, n=51 AOI from n=3 “Non-viral” patients and n=46 AOI from n=3 “Normal patients”. Data and metadata files (“expression_matrix.pq”, “metadata_matrix.pq”) were retrieved from <https://doi.org/10.5281/zenodo.4635286>, quantile normalised and log2(n+1) transformed prior to visualisation. (D) The boxplots show the expression of the selected genes in the bulk BAL alveolar macrophage RNA-seq data from (*3*) (n=72 COVID-19, n=5 Healthy Control, n=25 Non-Pneumonia Control, n=102 Other Pneumonia, n=28 Other Viral Pneumonia). Count data and metadata (Supplemental Data Files 2 and 3) were retrieved and the count data normalised with DESeq 2 and log2(n+1) transformed prior to plotting.

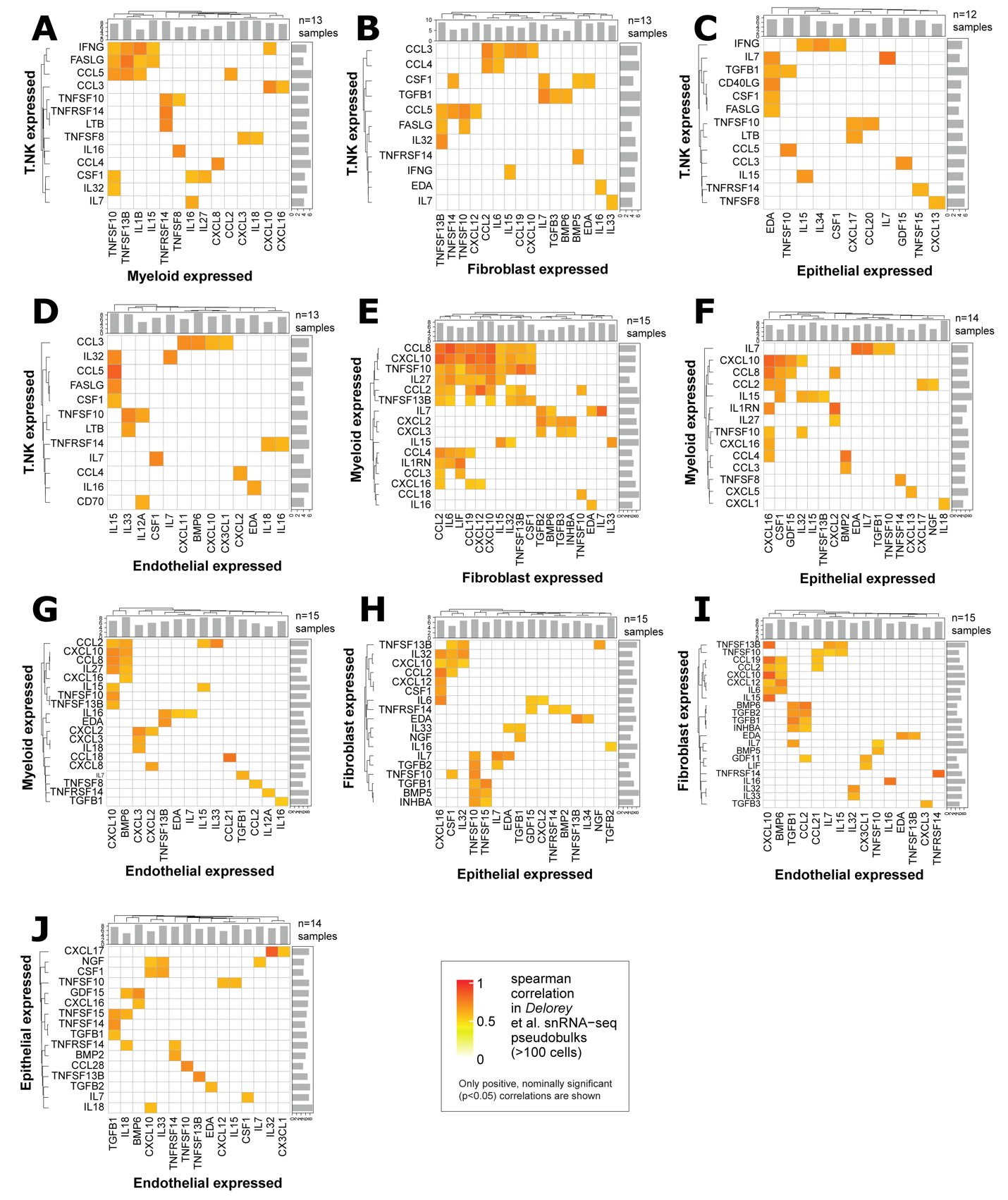

**Supplementary Figure 12: Analysis of correlation between cytokine expression in different cell types in COVID-19.** (A-J) The heatmaps show the correlations between cytokine expression in the indicated pairs of cell types across COVID-19 patients (number of patients retained for each cell type pairing is given in the top right). To avoid methodological bias, for this analysis only the nuclei data was included (n=17 samples from n=16 SARS-CoV-2 infected COVID-19 autopsy donors). Replicate samples from the same donor (D12) were merged. Pseudobulks of <100 cells were excluded. Summed pseudobulk counts were normalised within cell-type using DESeq2 prior to computation of correlation. Only positive, nominally significant (*p*<0.05) correlations are shown.

**Supplemental References**

1.	T. M. Delorey *et al.*, COVID-19 tissue atlases reveal SARS-CoV-2 pathology and cellular targets. *Nature*, (2021).

2.	D. J. Ahern *et al.*, A blood atlas of COVID-19 defines hallmarks of disease severity and specificity. *medRxiv*, 2021.2005.2011.21256877 (2021).

3.	R. A. Grant *et al.*, Circuits between infected macrophages and T cells in SARS-CoV-2 pneumonia. *Nature* **590**, 635-641 (2021).

4.	M. Liao *et al.*, Single-cell landscape of bronchoalveolar immune cells in patients with COVID-19. *Nat Med* **26**, 842-844 (2020).

5.	J. Schulte-Schrepping *et al.*, Severe COVID-19 Is Marked by a Dysregulated Myeloid Cell Compartment. *Cell* **182**, 1419-1440 e1423 (2020).

6.	K. J. Travaglini *et al.*, A molecular cell atlas of the human lung from single-cell RNA sequencing. *Nature* **587**, 619-625 (2020).

7.	A. F. Rendeiro *et al.*, The spatial landscape of lung pathology during COVID-19 progression. *Nature*, (2021).
